# Species-specific chromatin architecture and neurogenesis mediated by a human enhancer

**DOI:** 10.1101/2025.08.08.669395

**Authors:** Federica Mosti, Jing Liu, Katie Lam, Samantha Skavicus, Victoria A. Kapps, Ketrin Gjoni, Nicholas S. Heaton, Katherine S. Pollard, Debra L. Silver

## Abstract

Genomic modifications underlie the evolution of human features, including a larger neocortex and enhanced cognition. Human Accelerated Regions (HARs) are highly-conserved loci containing human-specific variants and can act as neurodevelopmental enhancers. However, the neurodevelopmental functions of HARs and their mechanism of gene regulation are largely unknown. Here, we show that human (*Hs*) *HAR1984* promotes neurogenesis by influencing species-specific transcription and chromatin interactions. *Hs-HAR1984* knock-in chimpanzee (*Pt*) cortical organoids contain more progenitors and neurons, whereas *Pt*-*HAR1984* knock-in human cortical organoids exhibit the opposite phenotype. *Hs-HAR1984* knock-in mice recapitulate increased neurogenesis, producing a thicker cortex with folding. Hi-C reveals *HAR1984* exhibits chromatin looping with its target genes *ETV5* and *TRA2B* in human fetal brains, notably reduced in chimpanzee, macaque and mouse neural cells. We show that human-specific mutations in *HAR1984* directly promote these interactions, favored by nearby structural variants. Further, we discover that human-specific ETV5 binding auto-regulates enhancer activity. This work demonstrates new molecular mechanisms underlying human-specific neurodevelopment, linking HARs to chromatin architecture, cortical cell fate and expansion and folding of brains.

## Introduction

The human brain is distinguished by its disproportionately large cerebral cortex relative to overall brain size. The cortex underlies many higher-order cognitive functions that set humans apart from other mammals, such as abstract reasoning, complex language, and flexible decision-making (Rilling, 2014; Sousa et al., 2017). The anatomy and function of the cortex is established during a tightly orchestrated process of corticogenesis, beginning with the transformation of neuroepithelial cells into apical radial glial cells (aRGCs) (Geschwind & Rakic, 2013; Lodato & Arlotta, 2015; Molnar et al., 2019; Silver, 2019). aRGCs generate glutamatergic neurons in an inside-out sequence, with deep-layer neurons formed first, followed by upper-layer neurons. These progenitors also produce neurogenic basal progenitors, including basal radial glia (bRGs) and intermediate progenitors (IPs), and eventually glia following neurogenesis (Cárdenas et al., 2018; Del-Valle-Anton et al., 2024; Pollen et al., 2015). The expansion of these basal progenitor populations is a hallmark of many larger-brained species, including humans (Borrell & Götz, 2014; Cárdenas et al., 2018; Del-Valle-Anton et al., 2024). This amplified progenitor pool contributes to a substantial increase in neurons and ultimately expanded neuroconnectivity (Huilgol et al., 2023; Preuss & Wise, 2022).

A critical question is how human-specific cellular neurodevelopmental features are encoded in the genome. Known uniquely human genomic elements include deletions, insertions and point mutations. Among the latter, Human Accelerated Regions (HARs) are DNA sequences that are evolutionarily conserved across mammals yet exhibit an unusually high rate of single-nucleotide substitutions in the human lineage (Bird et al., 2007; Bush & Lahn, 2008; Capra et al., 2013; Lindblad-Toh et al., 2011; Pollard et al., 2006; Prabhakar et al., 2006). Functional analyses demonstrate that approximately half of HARs act as regulatory transcriptional enhancers in neural cells, as indicated by epigenetic markers and massive parallel reporter assays (MPRAs) (Capra et al., 2013; Cui et al., 2025; Girskis et al., 2021; Uebbing et al., 2021; Won et al., 2019). Many HARs are located nearby genes involved in key neurodevelopmental processes such as neuronal proliferation, differentiation, and axon guidance (Boyd et al., 2015; Capra et al., 2013; Doan et al., 2016; Jorstad et al., 2023; Pollard et al., 2006; Prabhakar et al., 2006; Uebbing et al., 2021; Won et al., 2019). Recent work has demonstrated that HARs can directly influence cortical size and neuronal output by acting on neural progenitors (Boyd et al., 2015; Liu et al., 2025). In addition to neurodevelopment, HARs are implicated in other human traits, including limb development (Dutrow et al., 2022), male fertility (Norman et al., 2021), and thermoregulation (Aldea et al., 2021) as well as in disease susceptibility, including autism spectrum disorder (ASD), schizophrenia, Alzheimer’s disease, and cancer (Chen et al., 2018; Doan et al., 2016; Shin et al., 2024; Xu et al., 2015). Yet while there are >3000 HARs identified to date, known species-divergent functions for HARs are limited to these few examples.

The activity of enhancer elements is shaped by 3D genome architecture and chromatin dynamics. Within the nucleus, DNA is organized into a hierarchical structure, from chromosomal territories to topologically associating domains (TADs) and chromatin loops, which define the spatial relationships between regulatory elements and their target genes (Beagrie et al., 2017). TAD boundaries insulate regulatory interactions, creating modular units that facilitate precise gene regulation. During mouse neurodevelopment, structural rearrangements and chromatin remodeling permit context-specific activation of developmental programs and help define neural progenitor identity (Amberg et al., 2019; Harabula & Pombo, 2021). High throughput chromosome conformation capture (Hi-C) studies in human fetal brains and *in vitro* neural progenitor cells (NPCs) reveal that HARs physically interact with neurodevelopmental loci and thus may regulate them (Keough et al., 2023; Pal et al., 2025; Won et al., 2019). Comparative Hi-C analyses of human and chimpanzee NPCs show that HARs are frequently located within TADs containing human-specific structural variants (Keough et al., 2023). These variants are proposed to promote species-specific chromatin loops, which in turn influence enhancer-promoter interactions. Notably, HAR-enriched TADs also contain genes differentially expressed between human and chimpanzee brains, underscoring their role in species-specific regulatory architecture (Keough et al., 2023; Pal et al., 2025). However, direct links between HARs, chromosome architecture and neurogenesis have not been shown.

Here, we identified *HAR1984* as an enhancer that modulates human neurogenesis to impact brain anatomical features (Figure 1A). We used CRISPR-edited human and chimpanzee cortical organoids to demonstrate that *Hs-HAR1984* directly promotes neural progenitor proliferation and cortical cell identity to produce more neurons. To investigate this functional role *in vivo*, we generated a humanized *HAR1984* knock-in mouse model, which exhibits increased brain size and cortical folding, driven by expansion of progenitors and neurons. We identified *TRA2B* and *ETV5*, known neurodevelopmental regulators, as direct targets of *Hs-HAR1984*. We further demonstrated that ETV5, a transcription factor, binds *Hs-HAR1984* establishing a positive feedback loop to promote robust activation of *Hs-HAR1984*. Finally, we show that *Hs-HAR1984* promotes a human-specific chromatin loop with the promoters of *ETV5* and *TRA2B*, a less frequent topology found in non-human mammals. Modeling reveals this enhancer-promoter interaction is further stabilized by human-specific structural variants near *HAR1984*. Together, this provides a compelling example of how species-specific genomic architecture can contribute to human-specific neurodevelopmental traits.

**Figure 1.**
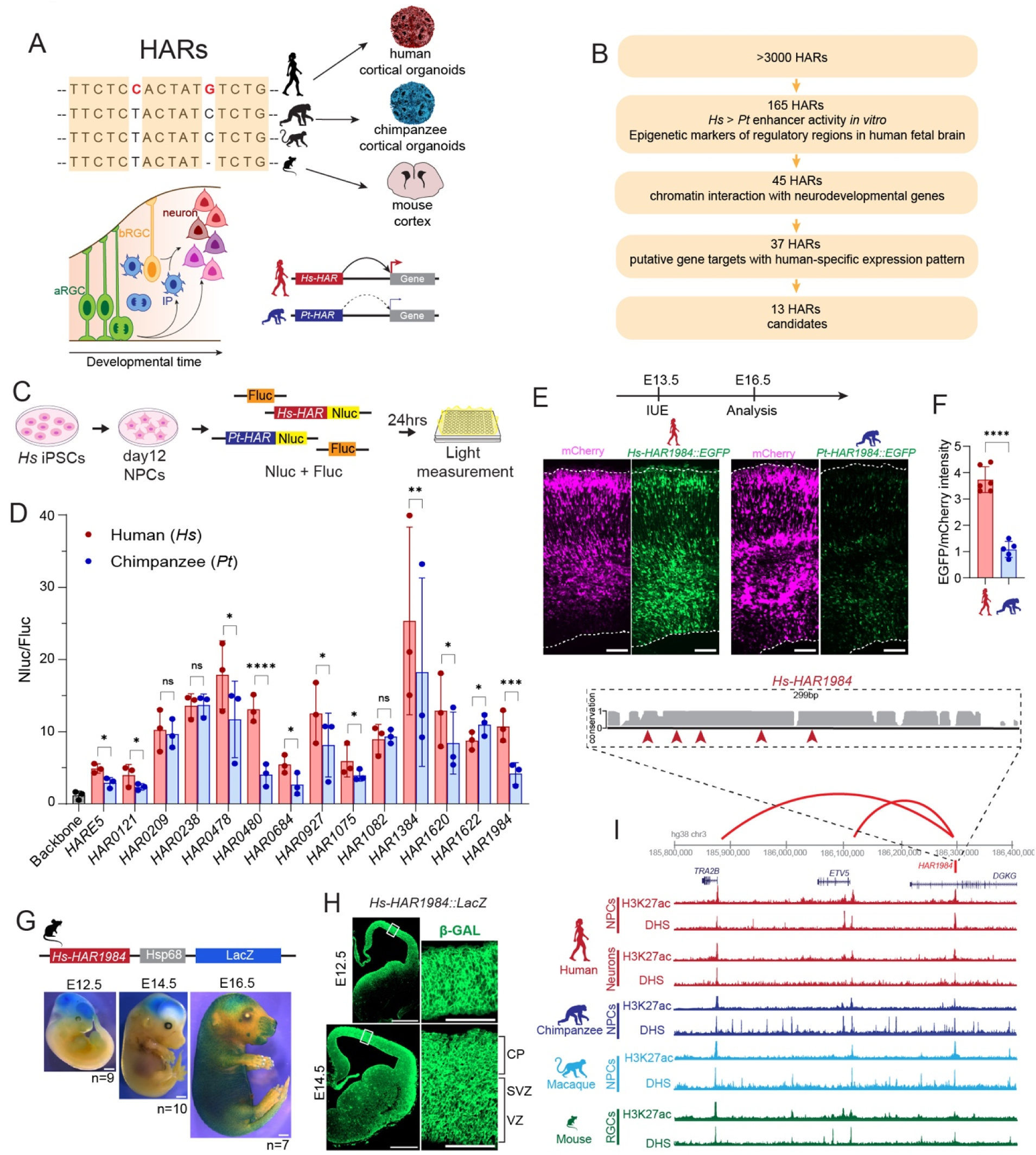
Identification of HARs with species-divergent enhancer activity during neurodevelopment. (A) A simplified representation of a HAR with conserved sequences (tan) and human-specific variants (red) (left). This work examines functions for HARs during human brain development using human, chimpanzee and mouse models (right). Neurogenesis cartoon (below) shows neurons produced by apical radial glial cells (aRGCs, green), intermediate progenitors (IPs, blue) and basal radial glial cells (bRGCs, yellow). (B) Experimental approach for *in silico* identification of neurodevelopmental HARs. (C) Experimental design for luciferase experiment. Human iPSCs were differentiated into neural progenitors (NPCs), plasmids containing Firefly luciferase (Fluc), and Nanoluciferase (Nluc) with human or chimpanzee HAR sequence were delivered at D12, and luciferase activity was assessed after 24hrs. (D) Nanoluciferase levels normalized to firefly control. Each dot represents independent differentiations. (E) *In utero* electroporation (IUE) paradigm with representative images. pCAGGS-mCherry (magenta) and *Hs-HAR1984::EGFP* or *Pt-HAR1984::EGFP* (green) were introduced at E13.5 and brains were analyzed at E16.5. (F) Quantification of intensity of EGFP relative to mCherry. Each dot represents independent embryos from 3 litters. (G) Sequence used to generate transgenic *Hs-HAR1984::LacZ* mice. Representative images of *Hs-HAR1984* β-gal activity at E12.5, E14.5 and E16.5. Numbers below indicate independent embryos analyzed from 2 independent transgenic lines. (H) β-gal immunofluorescence of *Hs-HAR1984::LacZ* cortices at E12.5 and E14.5. (I) *HAR1984* locus (top) with phastCons tracks showing locus conservation and location of 5 human-specific variants (red arrowheads). H3K27me3 ChIP-seq and ATAC-seq peaks (DHS, DNase Hypersensitivity) of *HAR1984* locus in human fetal cortex (NPCs and neurons), chimpanzee (NPCs), macaque fetal cortex (NPCs) and mouse (RGCs). *HAR1984* locus coordinates (hg38) and HiC interactions loops with *ETV5* and *TRA2B* from human fetal brain. Scale bars: 100 μm (E, H right), 500 μm (H left), 1mm (G). Two-way ANOVA with Bonferroni correction (D), Unpaired t-test (F). ns, not significant; \**p<*0.01; \*\**p<*0.001; \*\*\**p<*0.0001; \*\*\*\**p<*0.0001. Data are mean ± SD. See also Figure S1 and Table S1.

## Results

### Identification of Human Accelerated Regions with species-divergent enhancer activity during neurodevelopment

We first used *in silico* analysis to identify active enhancer HARs in neural progenitors with species-divergent regulatory activity between human (*Homo sapiens*, *Hs*) and chimpanzee (*Pan troglodytes*, *Pt*) sequences (Figure 1B). Amongst >3000 known HARs, we initially prioritized those with known species-divergent enhancer activity in neural cells. 165 were: a) located in open chromatin regions (Kundaje et al., 2015), b) enriched for H3K27ac and H3K4me1 (markers of active enhancer/promoters) in the human fetal brain (Won et al., 2016; Won et al., 2019), and c) showed stronger enhancer activity for the human compared to chimpanzee orthologue in neural cell lines (Girskis et al., 2021). We then focused on enhancers with known physical chromatin interactions with target genes implicated in neurodevelopmental function or expression. We defined 45 regions containing HARs interacting with almost 90 genes in human fetal brain or human neural progenitors (NPCs) (Jung et al., 2019; Won et al., 2019). For 37 of these HARs, at least one putative gene target showed human-specific temporal or spatial expression during brain development (Fietz et al., 2012; Florio et al., 2015; Florio et al., 2018; Johnson et al., 2015; Miller et al., 2014; Pollen et al., 2015). Amongst these, 13 HARs were selected as candidates for follow up analyses (Figure 1B, Table S1) (Please see methods for nomenclature used).

We next measured the relative enhancer activity of HAR orthologues in human NPCs. For this, we utilized luciferase assays in human NPCs derived from human induced pluripotent stem cells (iPSCs). We cloned each human and chimpanzee orthologue upstream of a nanoluciferase reporter with a minimal promoter and co-transfected these plasmids into NPCs with firefly luciferase reporter at day12 after neuronal induction. Light emission was measured after 24 hours (Figure 1C). All 13 of the human HAR sequences functioned as enhancers with at least 3 times more reporter expression than the minimal promoter alone (Figure 1D). The positive control (*HARE5*) (Boyd et al., 2015; Liu et al., 2025) as well as 10 HARs tested showed significant differential activity between species. For 9 HARs, the human sequence exhibited higher activity, similar to that reported in cell lines (Girskis et al., 2021), while 1 region (*HAR1622*) showed the opposite trend with the chimpanzee sequence exhibiting higher reporter levels. These data validate results of previous screens (Girskis et al., 2021; Uebbing et al., 2021) and nominate several candidates for further investigation.

We selected a subset of HARs (*HAR0480*, *HAR0684*, *HAR0927* and *HAR1984*) with pronounced divergent enhancer activity in human NPCs to assess if differential activity across species was also present *in vivo* during neurodevelopment. We employed *in utero* electroporation (IUE) to deliver an EGFP reporter plasmid driven by the human or chimpanzee HAR orthologue (*HAR::EGFP*), along with mCherry to report electroporated cells, in embryonic day (E) 13.5 mouse cortices. The electroporated embryos were then collected after 3 days (E16.5) for analysis (Figure S1A). All four HARs showed activity in the developing mouse cortex (Figure 1E-F, S1B-H), as reflected by robust EGFP levels across independent IUE’d brains. Notably, only two HARs (*HAR0480* and *HAR1984*) displayed significant differential activity between species *in vivo*, as quantified by the intensity ratio of EGFP/mCherry (Figure 1E-F, S1C, F). These data further demonstrate that HARs can act as enhancers, with evidence of species-divergent activity *in vivo*. Overall, our pipeline took advantage of *in silico*, *in vitro* and *in vivo* assays to successfully identify 10 regulatory regions with human-specific activity in NPCs, including 2 HARs with species differential activity in the mouse cortex.

### *Hs-HAR1984* shows robust activity throughout neurodevelopment

We focused in depth analysis on a particularly compelling HAR, *HAR1984* (*HANCS_612*) (Prabhakar et al., 2006). In our *in vitro* and *in vivo* screening, *HAR1984* was amongst the loci with the highest species-divergent enhancer activity in human NPCs ( ̴ 2.5 fold) and in the mouse cortex ( ̴3.5 fold) (Figure 1D-F). *HAR1984* is an intronic regulatory region of 299bp (hg38 chr3:186,239,308-186,239,606) containing 5 human-specific nucleotides (Figure 1I). This regulatory region displays peaks of H3K27ac and chromatin accessibility across the human fetal brain, chimpanzee NPCs and macaque and mouse embryonic cortex (Figure 1I) (Gorkin et al., 2020; Keough et al., 2023; Luo et al., 2021; Won et al., 2019). Moreover, in Hi-C datasets from human fetal brain and NPCs (Jung et al., 2019; Won et al., 2019), *HAR1984* establishes chromatin loops with two nearby genes, *ETV5* and *TRA2B*, suggesting possible regulatory functions (Figure S1I). *ETV5* (ETS variant transcription factor 5) is a member of the ETS transcription factor family and has been implicated in mouse neural progenitor differentiation and dendrite arborization (Fontanet et al., 2018). The splicing factor *TRA2B* (Transformer 2 Beta Homolog) is required in the mouse cortex for proper brain size, progenitor proliferation and survival (Roberts et al., 2014; Storbeck et al., 2014). In contrast, only weak interactions were observed with the promoter of *DGKG*, the gene containing *HAR1984* (Figure S1I). As *DGKG* is only expressed at low levels during human cortex development (Nowakowski et al., 2017), this suggests it is not a regulatory target of *HAR1984* during neurogenesis. This nominates two putative *HAR1984* regulatory targets, *TRA2B* and *ETV5*, during neurogenesis.

We next sought to understand *Hs-HAR1984* activity *in vivo*. Our IUEs suggest that *Hs-HAR1984* is active in the ventricular zone (VZ), subventricular zone (SVZ) and cortical plate (CP), based on GFP distribution across the developing cortex (Figure 1E). To further characterize the pattern of *Hs-HAR1984* activity *in vivo*, we generated a transgenic mouse in which LacZ reports *Hs-HAR1984* enhancer activity (*Hs-HAR1984::LacZ,* two independent transgenic lines analyzed). We observed robust LacZ activity in the developing mouse brain at E12.5, E14.5 and E16.5 (Figure 1G). At E12.5 and E14.5, *Hs-HAR1984* was almost exclusively active in the brain, particularly in the forebrain. At E16.5, *Hs-HAR1984* enhancer activity was additionally observed in the skin and whisker hair follicles. Coronal sections of E12.5 and E14.5 cortices showed β-gal immunostaining throughout all progenitor and neuron layers, similar to the IUE assay (Figure 1H). Thus, *Hs-HAR1984* is active *in vivo* in the developing cortex throughout key stages of neurogenesis.

### Genome-edited human and chimpanzee cortical organoids reveal *Hs-HAR1984* promotes neurogenesis

We next investigated the functional impact of *HAR1984* enhancer activity on human cortical neurogenesis. For this, we employed two complementary approaches using both human and chimpanzee cells. First, we used CRISPR-Cas9 genomic editing in human embryonic stem cells (ESCs) to replace *Hs-HAR1984* with *Pt-HAR1984*. We obtained two independent homozygous clones (*Hs-HAR1984^Pt/Pt^*). Second, we generated two independent *HAR1984* humanized chimpanzee iPSCs (*Pt-HAR1984^Hs/Hs^*) employing a similar genomic editing approach (Figure 2A). As controls for all subsequent experiments, we used a parental line and one non-edited clone each for human and chimpanzee cells. These CRISPR knock-in (KI) lines were then used to generate cortical organoids (Velasco et al., 2019). To validate cortical identity, each batch of differentiation was tested by qPCR for upregulation of forebrain markers, *PAX6* and *FOXG1,* and downregulation of iPSC marker, *OCT4,* at day 30 (D30) (Figure S2A-F). This revealed that all edited cell lines successfully differentiated into cortical fates, with no significant differences between lines.

**Figure 2.**
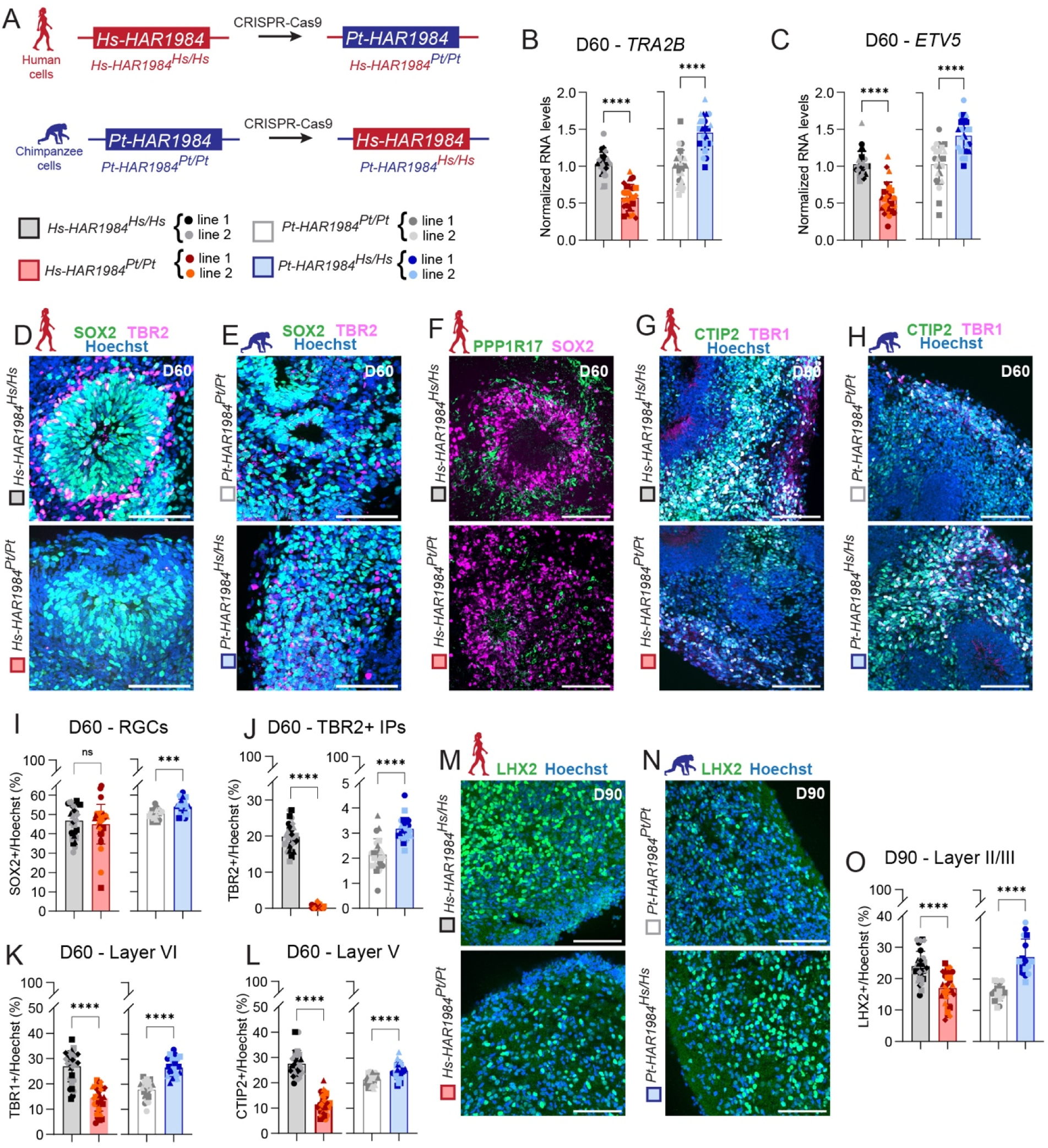
Genome-edited human and chimpanzee cortical organoids show *Hs-HAR1984* promotes neurogenesis. (A) Scheme of CRISPR-Cas9 editing in human ES and chimpanzee iPS cells. Two control lines and two knock-in lines for each species were used in all experiments. (B-C) RT-qPCR of *TRA2B* (B) and *ETV5* (C) levels normalized to control in human and chimpanzee D60 organoids. (D-E) Immunostaining for SOX2 (RGCs, green) and TBR2 (IPs, magenta) in D60 human (D) and chimpanzee (E) organoids. (F) Immunostaining for SOX2 (RGCs, magenta) and PPP1R17 (IPs, green) in D60 human organoids. (G-H) Immunostaining for CTIP2 (layer V neurons, green) and TBR1 (layer VI neurons, magenta) in D60 human (G) and chimpanzee (H) organoids. (I-L) Quantification of SOX2 (I), TBR2 (J), TBR1 (K), and CTIP2 (L) percentage over total number of cells (Hoechst) in D60 human and chimpanzee organoids shown in D-H. (M, N) Immunostaining for LHX2 (layer II/III neurons, green) in D90 human (M) and chimpanzee (N) organoids. (O) Quantification of LHX2 (O) percentage over total number of cells (Hoechst) in D90 human and chimpanzee organoids shown in M, N. Scale bars: 100 μm. Each dot represents a single organoid. Different dot shapes represent different batches of differentiation. D60 human organoids: 4 differentiations. D60 chimpanzee organoids and D90 human organoids: 3 differentiations. D90 chimpanzee organoids: 2 differentiations. 3-4 organoids were analyzed in each differentiation. Unpaired t-test. ns, not significant; \*\*\**p<*0.0001; \*\*\*\**p<*0.0001. Data are mean ± SD. See also Figure S2.

We next measured the expression levels of the *HAR1984* target genes, *ETV5* and *TRA2B*, in control and KI cortical organoids. At early stages of cortical development (D30), neither gene was differentially expressed in both human and chimpanzee KI organoids (Figure S2I-J). In contrast, at both D60 and D90, both genes were significantly reduced in *Hs-HAR1984^Pt/Pt^* and significantly increased in *Pt-HAR1984^Hs/Hs^*, relative to their respective controls (Figure 2B-C, S2P-Q). *DGKG*, the gene containing *HAR1984*, is expressed at low levels in cortical organoids (Uzquiano et al., 2022) and was unchanged in D60 KI organoids (Figure S2O). These data show that *Hs-HAR1984* is a transcriptional enhancer that is both necessary and sufficient for proper expression of *ETV5* and *TRA2B.* Further, it highlights roles for *HAR1984* in later stages (D60 and D90) of human and chimpanzee organoids.

We next used these human and chimpanzee organoids to assess neurogenesis by quantifying progenitors. We first assessed D30 when organoids are primarily composed of RGCs (Uzquiano et al., 2022). At this stage, the number of RGCs (SOX2+) and mitotic RGCs (SOX2+ PH3+) did not change in either human or chimpanzee KI organoids compared to controls (Figure S2G-H, K-L). This is consistent with the lack of significant changes in expression of *HAR1984* targets, *ETV5* and *TRA2B,* at D30. We next assessed progenitors at later stages in both KI organoids. Compared to human control, *Hs-HAR1984^Pt/Pt^* organoids showed a similar number of RGCs at D60, and a slight but significant decrease at D90 (Figure 2D, I, Figure S2M, T). *Pt-HAR1984^Hs/Hs^* organoids at D60 displayed slightly but significantly more RGCs with respect to chimpanzee control, with more a pronounced increase at D90 (Figure 2E, I, S2N, T). Thus, with elevated *HAR1984* enhancer activity (*Pt-HAR1984^Hs/Hs^)* we observe slight increases in RGCs, whereas the opposite is true with decreased activity (*Hs-HAR1984^Pt/Pt^*).

We also investigated IPs derived from RGCs by immunostaining. Strikingly, at both D60 and D90, compared to human control, *Hs-HAR1984^Pt/Pt^* organoids almost completely lacked expression for TBR2, a canonical marker of IPs (Figure 2D, J, S2M, U). To discriminate if this was due to loss of IPs or loss of TBR2 specifically, we assessed another IP marker, PPP1R17, in human organoids (Pebworth et al., 2021). We identified PPP1R17+ cells in *Hs-HAR1984^Pt/Pt^* confirming the presence of IPs (Figure 2F). Overall, we did not observe pyknotic nuclei around rosette regions, suggesting the absence of TBR2 expression is not due to selective cell death. Compared to chimpanzee control organoids, *Pt-HAR1984^Hs/Hs^* organoids at both D60 and D90 showed a significant increase of TBR2+ IPs (Figure 2E, J, S2N, U). These data show that higher *Hs-HAR1984* enhancer activity promotes IP numbers and may impact their identity. Taken together with the RGC analyses, this reveals an especially profound impact on TBR2+ IPs.

We next investigated how altered neurogenesis influences neuronal composition. Deep layer neurons are born earliest during cortical development and present in D60 organoids, while upper layer neurons appear at later stages by D90 (Lodato & Arlotta, 2015; Uzquiano et al., 2022). Relative to human controls, *Hs-HAR1984^Pt/Pt^* organoids at D60 and D90 exhibited a ∼50% reduction in deep layer neurons quantified using markers TBR1 (layer VI) and CTIP2 (layer V) (Figure 2G, K-L, S2R, V-W). In comparison, *Pt-HAR1984^Hs/Hs^* organoids showed significant increases for the same markers at D60 and D90, relative to their chimpanzee controls (Figure 2H, K-L, S2S, V-W). Focusing on upper layer neurons, we observed significantly fewer LHX2+ cells (layer II/III) in D90 *Hs-HAR1984^Pt/Pt^* organoids, compared to their respective human control (Figure 2M, O). In contrast, upper layer neurons were significantly increased in *Pt-HAR1984^Hs/Hs^* organoids, relative to chimpanzee controls (Figure 2N, O). These data indicate that *Hs-HAR1984* increases neuronal number across cortical layers. Taken all together, these cell biological analyses of progenitors in human and chimpanzee KI organoid models, demonstrate that *Hs-HAR1984* promotes neurogenesis in a species-specific fashion.

### Single cell transcriptomic analyses reveal *Hs-HAR1984* promotes cell fate specification in human cortical organoids

We next aimed to understand the impact of *Hs-HAR1984* on cell composition and cell fate at the transcriptional level. For this, we used single-cell RNA-seq (scRNAseq), focusing on the *Hs-HAR1984^Pt/Pt^* human organoids compared to their human control at both D60 and D90 (n=2 lines each genotype at each stage) (Figure 3A). After filtering the cells for quality, we obtained 65,506 cells (D60 controls: 15,406; D60 KI: 14,704 cells; D90 controls: 16,730; D90 KI: 18,716). Analysis of the whole dataset allowed us to obtain 37 Seurat clusters, from which we identified cell types using expression of established markers (Uzquiano et al., 2022; Velasco et al., 2019) (Figure 3B, S3A-B, D, Table S2). Differences in the relative proportion of progenitors and neurons were observed in both D60 and D90 organoids (Figure 3C).

**Figure 3.**
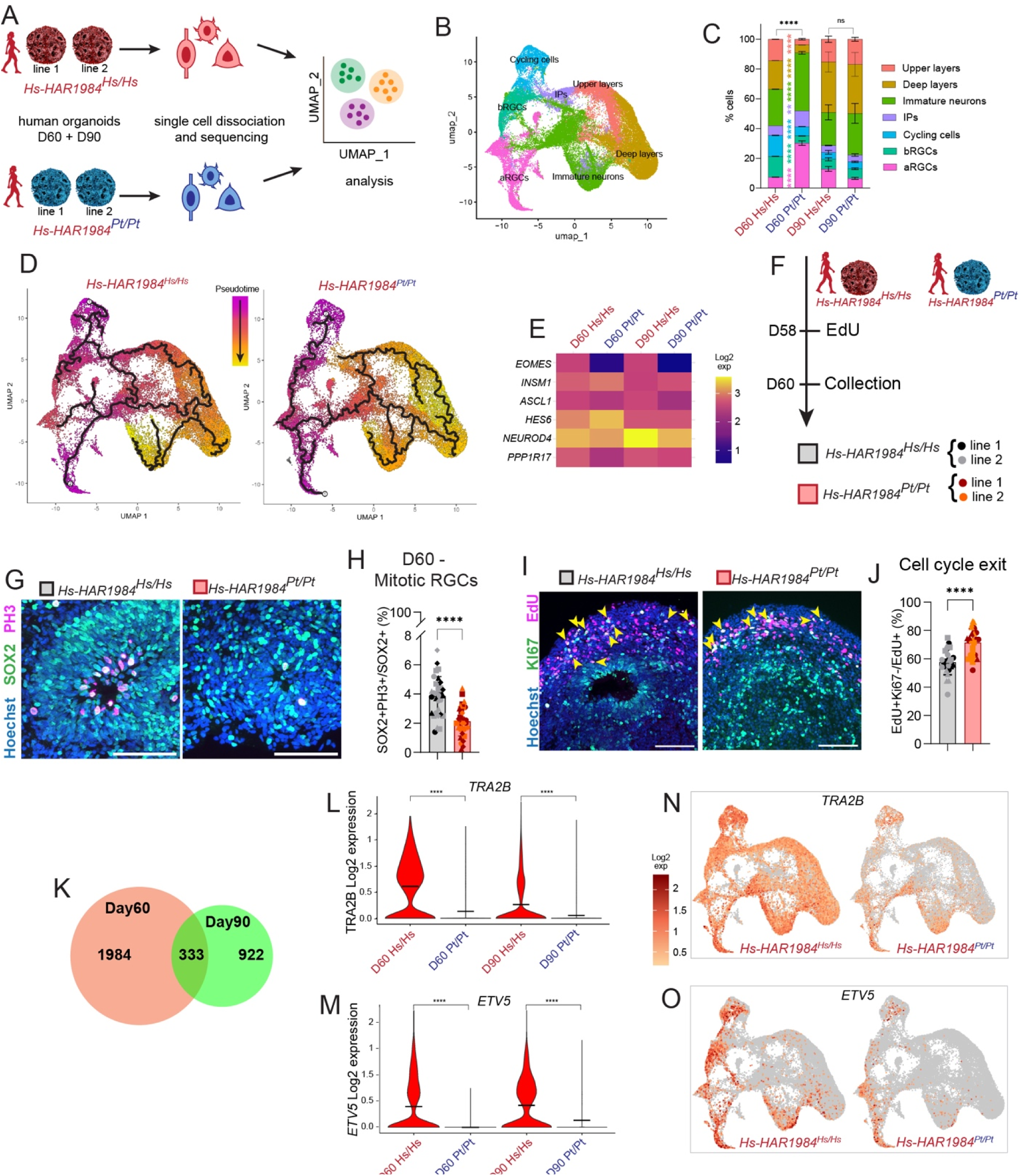
scRNAseq of human cortical organoids reveals *Hs-HAR1984* promotes cell fate during neurogenesis. (A) Scheme of experimental design. 2 control lines and 2 *Pt-HAR1984* knock-in line of human cortical organoids were dissociated and sequenced at D60 and D90. (B) UMAP showing integrated scRNAseq dataset labeled by cell identity. (C) Stack histogram of cell proportions between control and knock-in organoids at D60 and D90. (D) UMAP of cells colored by pseudotime trajectories: from less differentiated (magenta) to more differentiated (yellow). (E) Heatmap of gene expression levels of IP markers across all IP clusters. (F) Scheme of EdU pulse in organoids. EdU was added to the media of D58 human organoids and samples were collected after 48hrs. (G) Immunostaining for SOX2 (RGCs, green) and PH3 (mitotic cells, magenta) in D60 human organoids. (H) Quantification of SOX2+PH3+ percentage over total number of RGCs (SOX2) shown in G. 4 differentiations, 3-4 organoids per differentiations. (I) Immunostaining for KI67 (dividing cells, green) and EdU (magenta) in D60 human organoids. Arrows, KI67+EdU+ cells. (J) Quantification of EdU+KI67-(cells that exited cell cycle) over total EdU+ cells. 3 differentiations, 3-5 organoids per differentiation. (K) Venn diagram of DEGs from scRNAseq at D60 and D90. (L, M) *TRA2B* (L) and *ETV5* (M) expression from scRNAseq in human organoids at D60 and D90. (N, O) Feature plots of *TRA2B* (N) and *ETV5* (O) expression in integrated D60 and D90 dataset. Scale bars: 100 μm. Each dot represents a single organoid. Different dot shapes represent different batches of differentiation. Two-way ANOVA with Šídák correction (C). Unpaired t-test (H, J). Wilcoxon rank-sum test (L, N). ns, not significant; \*\*\*\**p<*0.0001. Data are mean ± SD. See also Figure S3-4 and Table S2.

We performed pseudotime analysis to bin cells according to their developmental state and model cell fate trajectories, dividing cells in 10 bins based on their differentiation state (Figure 3D). Over time, we observed a significant accumulation of more differentiated cells (bins 8-10) and a reduction of neural progenitors (bin 4) in *Hs-HAR1984^Pt/Pt^* compared to controls (Figure 3D, S4A). These data suggest possible alterations in progenitors and neurons which we next examined.

We first focused on RGC clusters. Compared to their control, D60 *Hs-HAR1984^Pt/Pt^* organoids contained proportionally more aRGCs (clusters 16, 19, 20, 23, 25, 26), but fewer bRGCs (clusters 2, 24, 33) (Figure 3B-C). Of note, in D60 *Hs-HAR1984^Pt/Pt^* organoids, subsets of aRGCs precociously expressed glial markers (clusters 16, 19, 26; *S100B*, *FGF1*, *CXCL14*) while others expressed ventral markers (clusters 20, 23; *DLX1, DLX5*) (Figure S3B-C, F). By D90, aRGC clusters expressing glial markers were enriched in control compared to *Hs-HAR1984^Pt/Pt^* (Figure S3D-E). Cycling progenitors were overall reduced in D60 *Hs-HAR1984^Pt/Pt^* organoids, and many of these clusters expressed RGC markers (clusters 17, 18, 28). Notably, while no difference was observed by immunofluorescence with SOX2+ RGCs in D60 *Hs-HAR1984^Pt/Pt^* organoids (Figure 2I), there were slight decreases by D90. Thus, the observed increase in transcriptionally-defined RGCs and decrease in cycling progenitors may reflect a precocious differentiation state of *Hs-HAR1984^Pt/Pt^* organoids.

We next investigated IPs in these single cell datasets. Our immunofluorescence analyses indicate that *Hs-HAR1984^Pt/Pt^* organoids have dramatically fewer TBR2+ IPs with expression of PPP1R17+IPs. Considering the specific downregulation of TBR2 (EOMES), we used several highly expressed IP markers (e.g., *ASCL1*, *INSM1*, *NEUROD4*) to assign cell type identity to this cluster (Martínez-Cerdeño et al., 2006; Pebworth et al., 2021) (Figure 3E, S3F). IPs were composed mainly by two clusters, one expressing *TBR2* (*EOMES)* (cluster 11) and one *TBR2*-negative (cluster 15). Notably, at D60, *TBR2+* cluster 11 were the main IPs in control organoids and markedly reduced in *Hs-HAR1984^Pt/Pt^*. In contrast the *TBR2-* cluster 15 was predominant in *Hs-HAR1984^Pt/Pt^* (Figure S3B-C, F, Table S2). This finding independently validates quantification of IPs by immunofluorescence and indicate that *Hs-HAR1984* is necessary for proper fate of IPs.

We next assessed the composition of neurons. Both upper and deep layer neuron clusters (cluster 0, 1, 4, 5, 7, 9, 10, 21) were reduced in *Hs-HAR1984^Pt/Pt^* compared to control at D60 (Figure S3B-C, F). This is similar to findings by immunostaining (Figure 2). In *Hs-HAR1984^Pt/Pt^*, immature neurons (cluster 3, 8, 12, 14, 27, 30) were significantly increased in D60 organoids, with some enriched at D90 (Figure S3D-E). These immature neuron clusters expressed some progenitor and interneuron markers, in addition to cortical markers (e.g., *SOX2*, *PAX6*, *DCX*, *CUX1*, *PDE1A*, *TBR1, DLX1, GAD2*) (Figure S3F Table S2). This suggests that *Hs-HAR1984* promotes proper neuronal differentiation and cell identity across development.

We next focused on the cycling cells to investigate progenitor proliferative capacity. Notably, cycling cell clusters (cluster 17, 18, 28, 29) were less represented in *Hs-HAR1984^Pt/Pt^* organoids compared to control at D60, suggesting a reduced progenitor proliferative state (Figure 3C, S3B-F). Of these only 1 cluster (29) was identified as IPs (Figure S3F). Using an established pipeline, we assigned cell cycle phases of G1/G0, S and G2/M to the progenitor clusters (Figure S4B) (Nestorowa et al., 2016). D60 *Hs-HAR1984^Pt/Pt^* organoids displayed significantly fewer cells in G2/M and S and an accumulation in G1/G0. Although cell cycle was not significantly different at D90, all phases showed similar trends as D60.

The G2/M reduction together with an increase in G0/G1 in KI organoids, suggests that *Hs-HAR1984* may promote proliferation of progenitors while tempering precocious cell cycle exit. Using immunofluorescence, we quantified PH3+SOX2+ cells in D60 *Hs-HAR1984^Pt/Pt^* organoids. This revealed a decrease in mitotic RGCs, validating the scRNA-seq analysis (Figure 3G-H). We also experimentally quantified the opposite phenotype in chimpanzee organoids with *Pt-HAR1984^Hs/Hs^* showing more mitotic RGCs at D60 (Figure S4C-D). These results indicate that *Hs-HAR1984* promotes RGC proliferation. We then assessed progenitor cell cycle exit by using EdU pulse-chase to label dividing cells and their progeny. We added EdU to the media of D58 human organoids and harvested them after 48hrs of incubation (Figure 3F). We co-stained organoids for the cell cycle markers KI67 and quantified the percentage of progenitors that exited the cell cycle (EdU+KI67-). Compared to human controls, we observed a significant increase in cell cycle exit in the *Hs-HAR1984^Pt/Pt^* organoids (control: 57.6%, KI:71.5%) (Figure 3I-J). This finding is consistent with the scRNA-seq data showing more immature neurons in *Hs-HAR1984^Pt/Pt^* organoids. Taken all together, these data indicate that *HAR1984* promotes neurogenesis by modulating progenitor proliferation and differentiation.

Finally, to understand how *HAR1984* impacts neurogenesis, we queried our dataset for differentially expressed transcripts between control and *Hs-HAR1984^Pt/Pt^* organoids. We identified 2,317 genes at D60 and 1,255 at D90 across clusters (fold change >1.5 and <-1.5, p-adj<0.05), with 333 transcripts in common between the two stages (Figure 3K). *HAR1984* target genes, *ETV5* and *TRA2B*, were both significantly reduced across timepoints (Figure 3L-O), corroborating observations by qPCR (Figure 2, S2). At both D60 and D90, *TRA2B* was ubiquitously expressed in all clusters, while *ETV5* was mostly expressed in aRGC and bRGCs at D60 with some expression in immature neurons at D90 (Table S2). Reduction in expression of both genes was observed across most clusters (Figure 3N-O, Table S2). This demonstrates that *Hs-HAR1984* increases expression of *ETV5* and *TRA2B* in multiple cell types across cortical neurogenesis. Gene Ontology (GO) analysis of the differentially expressed genes in common between D60 and D90, highlighted enrichment for genes related to neuronal differentiation and axonal guidance, as well as cerebral cortex regionalization and olfactory bulb interneurons (Figure S4E). *Hs-HAR1984^Pt/Pt^* lines displayed a downregulation of many cortical markers, while upregulating ventral telencephalon markers (Figure S3E-F, Table S2). Overall, these data suggest *Hs-HAR1984* contributes to robust expression of cortical markers to help promote cortical fate.

### *Hs-HAR1984* knock-in mouse model shows increased neurogenesis, brain size, and cortical folding

We next investigated functions of *Hs-HAR1984 in vivo*, by generating a KI mouse (*Mus Musculus*, *Mm*) model. We replaced *Mm-HAR1984* with *Hs-HAR1984* using iGONAD CRISPR-Cas9 methods (Gurumurthy et al., 2019) (Figure 4A). We obtained a *Hs-HAR1984* KI heterozygous mouse line (*Mm-HAR1984^Hs/Mm^*), giving us the opportunity to study the function of the human sequence of *HAR1984 in vivo*. To assess the impact of manipulating the *HAR1984* locus, we also used CRISPR to generate a *Mm-HAR1984* KO mouse line with either heterozygous or complete deletion (*Mm-HAR1984^del/Mm^* or *Mm-HAR1984^del/del^*) (Figure S5J). For both KI and KO models, the entire *HAR1984* locus and the homology arms (mm9 chr16:22363786-22364363) were sequenced, which confirmed expected gene editing. Adult heterozygous male *Mm-HAR1984^Hs/Mm^* mice showed reduced fertility, while neither *Mm-HAR1984^del/Mm^* male nor females showed any fertility issues. To understand the basis for the fertility reductions in *HAR1984* humanized mouse lines, we measured their testes (Figure S5A). Compared to control, *Mm-HAR1984^Hs/Mm^* exhibited significantly decreased testes weight after normalization to body weight (Figure S5B). The testes showed significant overexpression of *HAR1984* target genes, *Etv5* and *Tra2b* but not *Dgkg* (Figure S5C-E). Both *Etv5* and *Tra2b* are implicated in testes integrity and spermatogonia niche maintenance (Dalgliesh et al., 2025; Morrow et al., 2007; Tyagi et al., 2009; Zhang et al., 2021), suggesting that their overexpression in this tissue might be detrimental for fertility. *Mm-HAR1984^Hs/Mm^* females did not show fertility issues. These data show that *Hs-HAR1984* is active in multiple tissues, playing a role in male fertility, likely through upregulation of *Etv5* and *Tra2b*.

**Figure 4.**
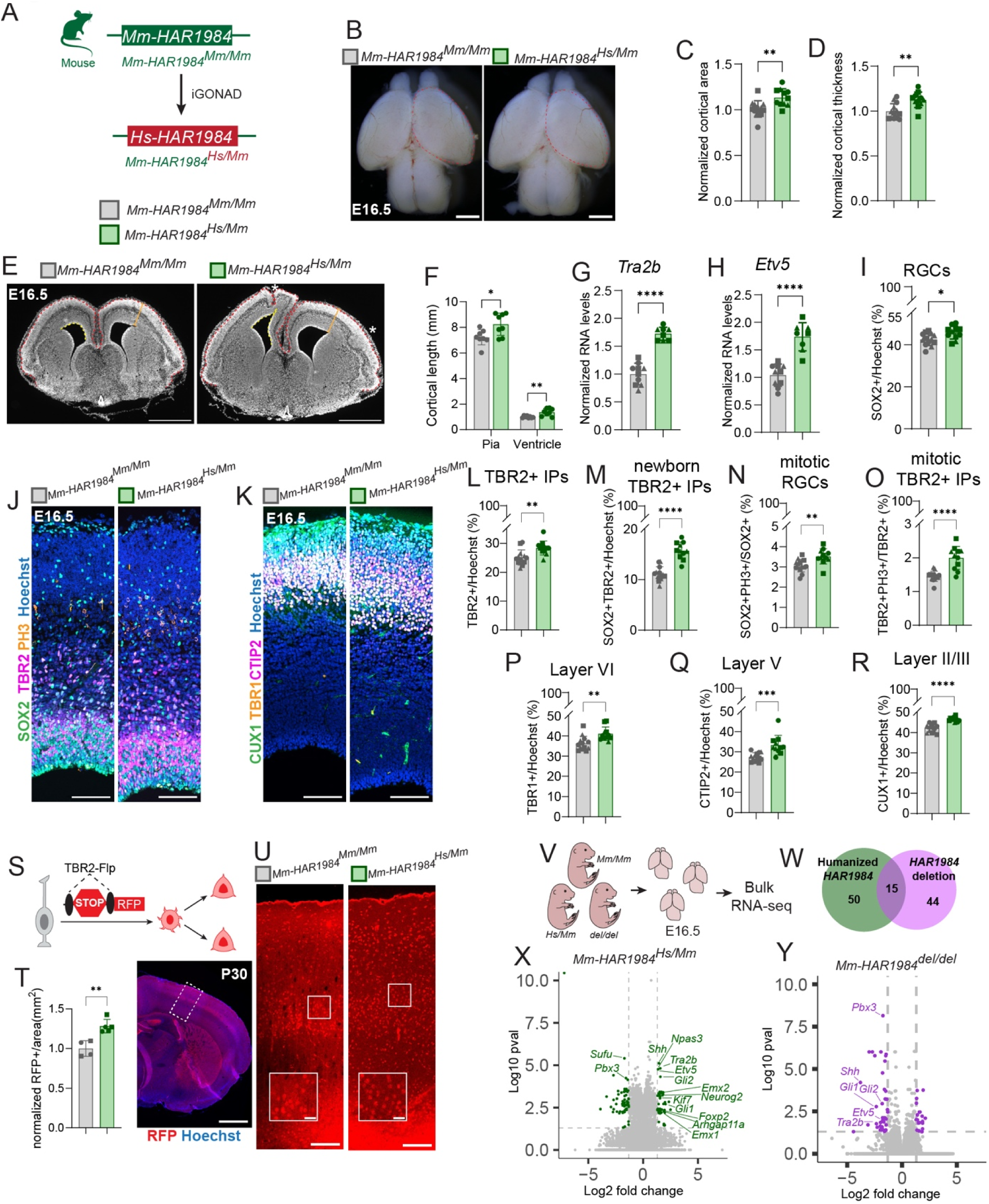
*Hs-HAR1984* knock-in mouse model shows increased neurogenesis, brain size and cortical folding. (A) Scheme of iGONAD knock-in to replace *Mm-HAR1984* with *Hs-HAR1984*. (B) Whole mount images of E16.5 control and humanized mouse brains. Red dotted line of control superimposed on knock-in brain. (C-D) Quantifications of E16.5 cortical area (C) and cortical thickness (D). (E) Coronal sections of E16.5 control and humanized brains. Red line is cortical length on pia side, yellow line is length on ventricle side, orange indicate cortical thickness. Asterisks indicate folded regions. (F) Measurement of cortical length on pia and ventricle side as indicated in E. (G-H) RT-qPCR of *Tra2b* (G) and *Etv5* (H) levels normalized on control at E16.5 across 3 litters, control n=10; humanized n=8. (I) Quantification of SOX2 percentage over total number of cells (Hoechst) shown in J. (J) Immunostaining for SOX2 (RGCs, green) and TBR2 (IPs, magenta) and PH3 (mitotic cells, orange) in E16.5 mouse cortices. (K) Immunostaining for TBR1 (layer VI neurons, orange), CTIP2 (layer V neurons, magenta) and CUX1 (layer II/III neurons, green) in E16.5 mouse cortices. (L-M) Quantification of TBR2 (L) and SOX2+TBR2+ (M) percentage over total number of cells (Hoechst) shown in J. (N-O) Quantification of SOX2+PH3+ percentage over total number of RGCs (SOX2) and TBR2+PH3+ percentage over total number of IPs (TBR2) shown in J. (P-R) Quantification of TBR1 (P) CTIP2 (Q) and CUX1 (R) percentage over total number of cells (Hoechst) shown in K. (S) Overview of lineage tracing of indirect neurogenesis in P30 cortex (T) Quantification of RFP^+^ cells in P30 control (*n* =4) and *Hs-HAR1984^Hs^*^/*Hs*^ (*n* =5) cortices (left). P30 genetically labelled by RFP (red) and DAPI (blue) (right). (U) P30 control and *Mm-HAR1984^Hs^*^/*Mm*^ stained for RFP (red) and DAPI (blue) with an enlarged image (inset) (V) Bulk RNA-seq experimental design. Three cortices each from 2 independent litters for control, *Mm-HAR1984^Hs/Mm^* and *Mm-HAR1984^del/del^* were sequenced. (W) Venn diagram of DEGs from bulk RNA-seq of *Mm-HAR1984^Hs/Mm^* and *Mm-HAR1984^del/del^* cortices. (X, Y) Volcano plot of genes from bulk RNA-seq of *Mm-HAR1984^Hs/Mm^* mouse (X) and *Mm-HAR1984^del/del^* mouse (Y) normalized on control. Green (X) and purple (Y) dots are significant differentially regulated genes; grey dots are not significant. Biological meaningful hits are highlighted. Scale bars: 20 μm (U inset), 100 μm (J-K, U), 500 μm (E), 1mm (B, T). Each dot represents a single embryo. Different dot shapes represent independent litters (N=3), control n=13; humanized n=11 (C-D, I, L-O, P-R). Unpaired t-test. ns, not significant; \**p<*0.01; \*\**p<*0.001; \*\*\**p<*0.0001; \*\*\*\**p<*0.0001. Data are mean ± SD. See also Figure S5 and Table S3.

We next focused on the role of *Hs-HAR1984* for cortical development *in vivo*. We first measured cortical size and cortical features of mouse KI brains. *Mm-HAR1984^Hs/Mm^* adult cortices were significantly enlarged and increased 9% in weight compared to control brains, after normalization to body weight (Figure S5F-G). We next analyzed brain size at E16.5, a stage when *Hs-HAR1984* is transcriptionally active (Figure 1). We quantified a slight but significant 13% increase in cortical area of *Mm-HAR1984^Hs/Mm^* compared to control (Figure 4B-C). *Mm-HAR1984^Hs/Mm^* cortices were also significantly thicker than control (Figure 4D-E). Notably, 43% of *Mm-HAR1984^Hs/Mm^* brains showed patterns of cortical folding (9/21 *HAR1984^Hs/Mm^* from 5 independent litters) (Figure 4E, S5I). Folding was variable across independent animals, evident in lateral and medial regions, as well as different rostro-caudal axes (Figure S5I). This is striking as mice are lissencephalic and indicates that *Hs-HAR1984* may contribute to cortical folding in humans. We further measured cortical length at both pial and ventricular surfaces, quantifying a 13% and 25% increase, respectively, in *HAR1984^Hs/Mm^* mice compared to control (Figure 4E-F). In contrast to the KI, heterozygous and homozygous deletion of *Mm-HAR1984* did not impact brain area or cortical thickness at E16.5 (Figure S5K-M). This suggests that the phenotypes observed in *Mm-HAR1984^Hs/Mm^* are driven by *Hs-HAR1984* and not by a modification of the genomic locus. Overall, these data demonstrate that activity of *Hs-HAR1984* during embryonic stages significantly affects mouse brain size and anatomy, associated with a thicker and tangentially longer cortex and presence of cortical folding.

We investigated if these anatomical changes in *Mm-HAR1984^Hs/Mm^* brains were accompanied by molecular alterations in target genes. At E16.5, expression of both *HAR1984* target genes, *Tra2b* and *Etv5*, was significantly increased in *Mm-HAR1984^Hs/Mm^* cortices compared to control (fold change ̴1.7 for both genes) (Figure 4G-H). This corroborates similar increases in target gene expression in humanized D60 KI chimpanzee organoids (Figure 2). In comparison, E16.5 heterozygous or homozygous deletion of *Mm-HAR1984* caused a decrease of both *Etv5* and *Tra2b*, relative to the control (Figure S5N-O). Similar to *Hs-HAR1984* KI in chimpanzee organoids and mice, *Dgkg* expression was unaffected in either humanized or deletion mouse model (Figure S5H, P). These analyses of KI and KO brains indicate that *Hs-HAR1984* controls expression of its target genes in the developing cortex, supporting the validity of this model. Taken together with organoid data, *Hs-HAR1984* shows increased enhancer activity to regulate its target genes, regardless of the *trans* environment (chimpanzee or mouse).

In order to understand the mechanisms underlying increased cortical size, we examined neural progenitor populations in the KI mice. At E16.5, *Mm-HAR1984^Hs/Mm^* cortices contained slight but significant increases in RGCs relative to control (Figure 4I-J). Likewise, we quantified significantly more total and newborn IPs (SOX2+TBR2+) in *Mm-HAR1984^Hs/Mm^* cortices, relative to control (Figure 4L, M). Both RGCs and IPs showed increased proliferation in *Mm-HAR1984^Hs/Mm^* cortices, relative to control, as assessed by PH3 staining (Figure 4N-O). Importantly, these data corroborate findings in humanized D60 KI chimpanzee organoids, in which introduction of *Hs-HAR1984* increased both RGC and IP number as well as RGC proliferation (Figures 2, S4D). These data demonstrate that *Hs-HAR1984* promotes neural progenitor proliferation leading to expansion of both RGC and IP populations, with an especially notable increase of IPs.

We then measured the impact of *Hs-HAR1984* upon cortical excitatory neurons at E16.5. Layer VI and V neurons, marked by TBR1 and CTIP2, respectively, both showed slight but significant increases in *Mm-HAR1984^Hs/Mm^* cortices compared to control (Figure 4K, P-Q). Similarly, relative to control littermates, upper layer neurons were also significantly increased in *Mm-HAR1984^Hs/Mm^* (Figure 4K, R). At E16.5 these neurons are still undergoing neurogenesis. This increase in neurons spanning deep and upper layers phenocopies humanized *HAR1984* KI chimpanzee cortical organoids. Taken together, these data demonstrate that *Hs-HAR1984* promotes progenitor proliferation which leads to increased neuron number and ultimately enlarged cortices.

*Hs-HAR1984* has a profound effect in TBR2+ IPs in both E16.5 mouse embryos and human and chimpanzee cortical organoids. Given this, we next investigated the fate of IPs in adult mice to determine if this progenitor expansion resulted in more progeny. We used an Ai65-FRT reporter under the control of *TBR2*-FLP to label IP-derived progeny in postnatal day 30 (P30) mouse cortex (Huilgol et al., 2023) (Fig. 4S). *Mm-HAR1984^Hs^*^/*Mm*^ cortices, RFP-labelled progeny significantly increased relative to the control (fold change ̴1.3) (Fig. 4T-U). This data suggest that *Hs-HAR1984* promotes neuron production by increasing indirect neurogenesis through IP expansion.

### *Hs-HAR1984* is necessary and sufficient for expression of key pathways controlling corticogenesis

We next aimed to understand the molecular mechanisms by which *Hs-HAR1984* affects cortical neurogenesis. Towards this we performed bulk RNA-seq on E16.5 cortices from *Mm-HAR1984^Hs/Mm^*, *Mm-HAR1984^del/del^*, and their control littermates. Our goal was to identify downstream affected transcripts with humanized *HAR1984* and *HAR1984* deletion (Figure 4V). We identified 65 differentially regulated transcripts for *Mm-HAR1984^Hs/Mm^* and 59 for *Mm-HAR1984 ^del/del^* relative to their respective littermate controls (fold change >1.3 and <-1.3, p-value<0.05) (Table S3). Of these, 15 transcripts were dysregulated in both humanized and deletion *HAR1984* cortices (Figure 4W, Table S3). The shared transcripts included *HAR1984* targets, *Etv5* and *Tra2b*, showing expected opposite trends in humanized (upregulated) and deletion (downregulated) mice (Figure 4X-Y). This is in accordance with what we previously observed by RT-qPCR (Figure 4G-H, Figure S5N-O).

Amongst the other common dysregulated transcripts were components of the Sonic hedgehog (Shh) pathway, a regulator of brain development (Cai et al., 2023). *Mm-HAR1984^Hs/Mm^* samples showed significant upregulation of Shh pathway activators (*Shh*, *Gli1*, *Gli2* and *Kif7*) and downregulation of inhibitor components (*Sufu*). In comparison, deletion of *HAR1984* led to downregulation of these Shh pathway components (*Shh*, *Gli1*, *Gli2*). We independently validated these alterations of *Shh*, *Gli1* and *Gli2* by RT-qPCR of *Mm-HAR1984^Hs/Mm^* and *Mm-HAR1984^del/del^* cortices (Figure S5Q-S). Increased and decreased activation of Shh pathway in the cortex is associated with increased cortical folding and brain size in mouse and ferrets (Hou et al., 2021; Komada et al., 2008; Wang et al., 2016; Yabut et al., 2015), suggesting possible molecular contributions for cortical folding in *HAR1984* mouse model.

To further understand *HAR1984* function, we focused on 29 transcripts upregulated solely in *Mm-HAR1984^Hs/Mm^* samples compared to control. Amongst these, we observed enrichment for known cortical neural progenitor and neuronal regulators (*Arhgap11a*, *Emx1*, *Emx2*, *Foxp2*, *Neurog2*, *Npas3*) (Cecchi & Boncinelli, 2000; Co et al., 2020; Lu et al., 2023) (Figure 4X, Table S3). Of note, these same transcripts were significantly reduced in scRNA-seq of *Hs-HAR1984^Pt/Pt^* human cortical organoids (Table S2). These findings, from both *in vivo* and *in vitro* mouse and human models, suggest that *Hs-HAR1984* facilitates neurogenesis by promoting proper expression of key fate determinants.

### *TRA2B* and *ETV5* knock-down phenocopies *HAR1984*, and ETV5 generates a positive feedback loop

Our organoid and mouse studies point to *ETV5* and *TRA2B* as major targets of *HAR1984.* Given this, we probed our scRNA-seq datasets to determine if targets of ETV5 transcription factor or TRA2B RNA binding protein were also impacted by the presence of *Hs-HAR1984^Pt/Pt^* (Anderson et al., 2012; Best et al., 2014; Grellscheid et al., 2011; Kalkan et al., 2019; Liu & Zhang, 2019; Storbeck et al., 2014; Uren et al., 2012). In *Hs-HAR1984^Pt/Pt^* organoids at both D60 and D90, known ETV5 and TRA2B targets were significantly dysregulated (Figure S6A-B, Table S2). Given this, we tested if modulating *ETV5* and *TRA2B* levels phenocopies *Hs-HAR1984* impact on gene expression. We manipulated *ETV5* and *TRA2B* expression using dCas9-KRAB to epigenetically inhibit their promoters, and used a similar approach to independently inhibit the *HAR1984* enhancer. First, using a human iPSC line which constitutively expresses BFP-dCas9-KRAB, we generated cortical organoids. At D55 we dissociated cells and delivered sgRNA targeting either *ETV5* promoter, *TRA2B* promoter, *HAR1984,* or a mock sequence (control) (Figure 5A). 5 days after transduction we collected cells for RNA extraction and immunostaining (Figure 5B-E). RT-qPCR confirmed significant downregulation of *ETV5* and *TRA2B* following independent targeting of either promoter or simultaneous targeting, relative to the mock control (Figure 5C-D). Moreover, *TRA2B* levels were significantly reduced after inhibition of the *ETV5* promoter (Figure 5C). This indicates the ETV5 transcription factor also regulates *TRA2B* expression. Consistent with its role as a transcriptional enhancer, epigenetic silencing of *Hs-HAR1984* also reduced *TRA2B* and *ETV5* (Figure 5C-D).

**Figure 5.**
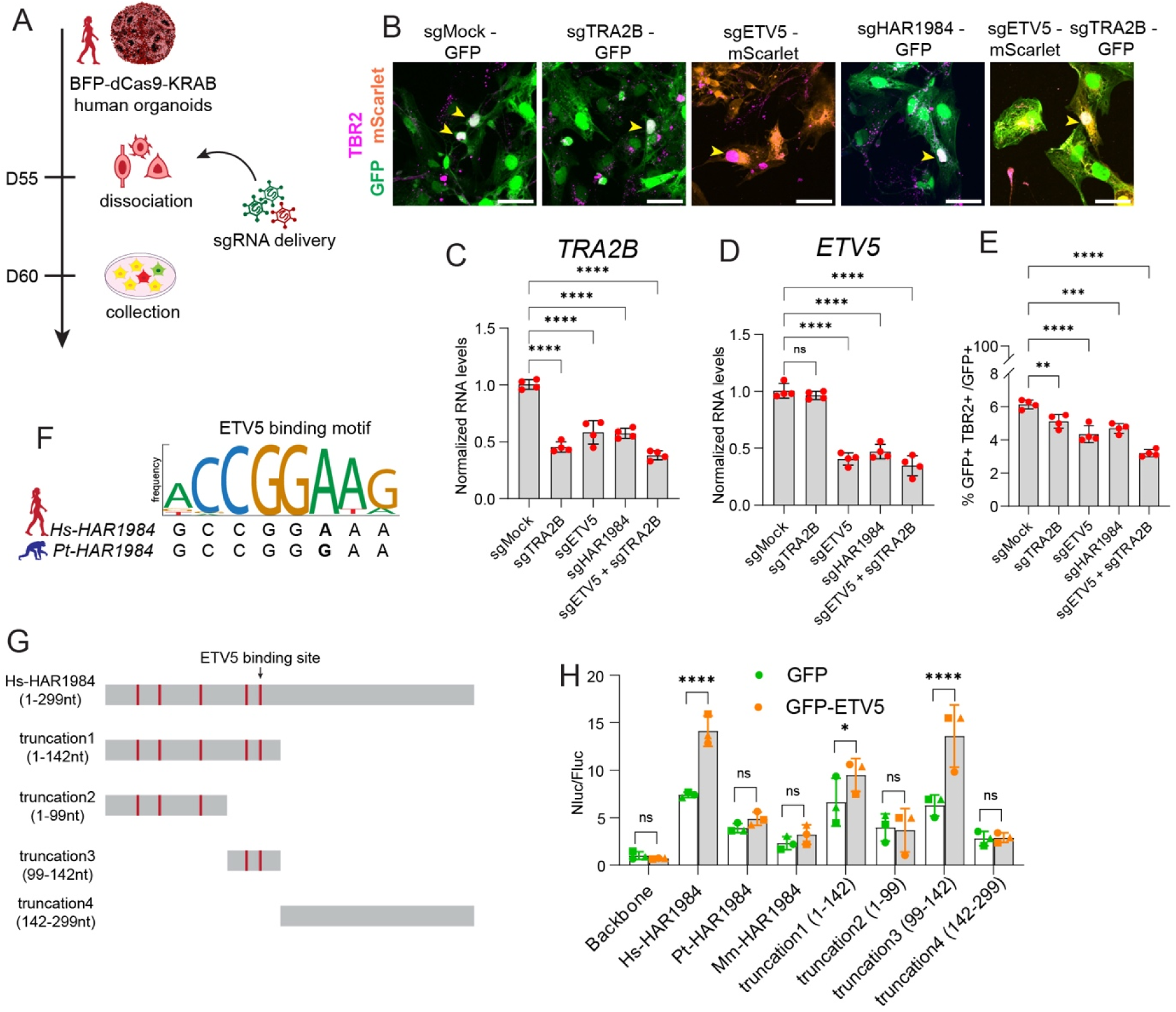
*TRA2B* and *ETV5* knock-down phenocopies *HAR1984*, and ETV5 generates a positive feedback loop. (A) Experimental design for epigenetic manipulation in human cells. BFP-dCas9-KRAB iPSCs were used to generated human cortical organoids. At D55 organoids were dissociated and sgRNA for *ETV5* and *TRA2B* promoters, *HAR1984* or mock were delivered. Cells were collected after 5 days *in vitro*. (B) Representative images from dissociated organoids after 5 days of sgRNAs (green or orange) viral transduction and immunostained for TBR2 (magenta). (C-D) RT-qPCR expression for *TRA2B* and *ETV5* in dissociated organoids 5 days after receiving sgRNAs. (E) Quantifications of GFP+TBR2+ cells normalized over GFP+ cells shown in B. (F) Predicted ETV5 binding motif on *Hs-* and *Pt-HAR1984* sequences with human-specific variant (bold). Size of letters is proportional to frequency of nucleotide in that position in consensus sequence. (G) Scheme of *Hs-HAR1984* human-specific variants (red) and constructs used in luciferase assay shown in H. (H) Nanoluciferase (Nluc) levels normalized to firefly (Fluc) control in D12 NPCs. GFP or GFP-ETV5 were co-delivered with luciferase reporter. Each dot represents an independent differentiation batch (n=3 technical replicates for each differentiation). Scale bars: 10 μm. Each dot represents a single differentiation. Two-way ANOVA (C-E, H). ns, not significant; \**p<*0.01; \*\**p<*0.001; \*\*\**p<*0.0001; \*\*\*\**p<*0.0001. Data are mean ± SD. See also Figure S6 and Table S4.

We next assessed the phenotypic impact of reducing *HAR1984* target gene expression as well as independently silencing *HAR1984*. For this we assessed IPs following epigenetic inhibition. Using *sgHAR1984* we observed a significant 24% reduction of TBR2+ IPs. This phenocopied loss of IPs seen in D60 *Hs-HAR1984^Pt/Pt^ KI* organoids, which also have reduced enhancer activity and target gene expression (Figure 2B-C). Similarly, targeting either *TRA2B* or *ETV5* also significantly reduced IPs by ∼17% and 29%, respectively, and targeting both genes simultaneously reduced TBR2+ cells by half (Figure 5B, E). In contrast, the number of RGCs was not affected with any of the sgRNA conditions (Figure S5C-D). This is consistent with the relatively mild impact on RGCs in *Hs-HAR1984^Pt/Pt^ KI* organoids (Figure 2I). Overall, these data demonstrate that downregulation of *TRA2B* and *ETV5* phenocopies *Hs-HAR1984* role in production of TBR2+ IPs. Taken together with Hi-C data, this further indicates that both genes are the main *Hs-HAR1984* targets contributing to neurogenesis.

We next aimed to understand how the human-specific *HAR1984* sequence influences enhancer activity. To test this, we performed a nanoluciferase assay in human NPCs as previously (Figure 1C). We tested *Hs-*, *Pt-* or *Mm-HAR1984* as well as 4 truncations of *Hs-HAR1984* containing all human specific sites (1-142 nt), the first three mutations (1-99nt), the last two mutations (99-142nt), or the conserved C-terminus with no human specific sites (142-299nt) (Figure 5G). As expected, relative to *Hs-HAR1984, Pt-HAR1984* had significantly tempered luciferase activity, as did *Mm-HAR1984* (Figure S6E). Truncations which removed the last two *Hs*-specific variants or all five (truncations 2 and 4) significantly reduced luciferase activity (Figure S6E). These data reinforce the importance of the *Hs*-specific variants, especially the last two, in promoting enhancer activity.

We next used transcription binding site predictions on *Hs-HAR1984* and *Pt-HAR1984* to understand trans-factors important for this species-specific enhancer activity (Martin et al., 2019). We identified multiple transcription factor families that were differentially affected across species (Table S4), either losing or gaining a binding site in *HAR1984* in humans. One of these was the ETS transcription factor family, which gained a new binding site in the human *HAR1984* sequence due to the last human-specific variant (Figure 5F). This is notable as ETV5 is a member of the ETS transcription factor family (Wei et al., 2023). Moreover, our CRISPR epigenetic manipulating data indicate that *ETV5* downregulation reduces *TRA2B* gene expression in cortical organoids (Figure 5C). Given this, we postulated that ETV5 may have an amplifying impact on *HAR1984* enhancer activity by trans-activation and binding. To test this possibility, we measured the extent to which ETV5-EGFP modifies *HAR1984* enhancer activity using the luciferase assay. ETV5-EGFP overexpression specifically increased enhancer activity of full-length *Hs-HAR1984.* In contrast, it had no impact on enhancer activity of either chimpanzee or mouse *HAR1984* (Figure 5H). ETV5-EGFP significantly amplified *HAR1984* enhancer activity only for those sequences containing the predicted human-specific binding site (truncations 1 and 3). These data demonstrate that *Hs-HAR1984* gained a human specific ETV5 binding site and that ETV5 binds its own enhancer, *Hs-HAR1984*. Together this creates a positive feedback loop of regulation to promote human neurogenesis.

### *Hs-HAR1984* locus promotes species-specific chromatin interactions with *ETV5* and *TRA2B*

Differences in enhancer regulation of specific target genes might be due to changes in nuclear architecture and chromatin loop dynamics. To elucidate the molecular mechanisms by which *HAR1984* regulates *ETV5* and *TRA2B*, we analyzed chromatin interactions in the *HAR1984* locus. Using Hi-C datasets from human, macaque and mouse embryonic brains and chimpanzee NPCs (Gorkin et al., 2020; Keough et al., 2023; Luo et al., 2021; Won et al., 2019), we investigated the frequencies of physical interactions between enhancers and putative target regions. This analysis revealed an expected enrichment in interactions between *HAR1984* and *ETV5* and *TRA2B* promoters in human fetal brains (Figure 6A). Notably, this interaction showed reduced frequency in macaque fetal brains(Figure 6B). Inspection of Hi-C data from chimpanzee NPCs (Figure 6K, bottom left) and mouse fetal brain indicate reduced frequency in other mammals as well (Gorkin et al., 2020; Keough et al., 2023). These data indicate that *HAR1984* exhibits chromosome looping to contact its target genes, with distinct strength of interactions across species.

**Figure 6.**
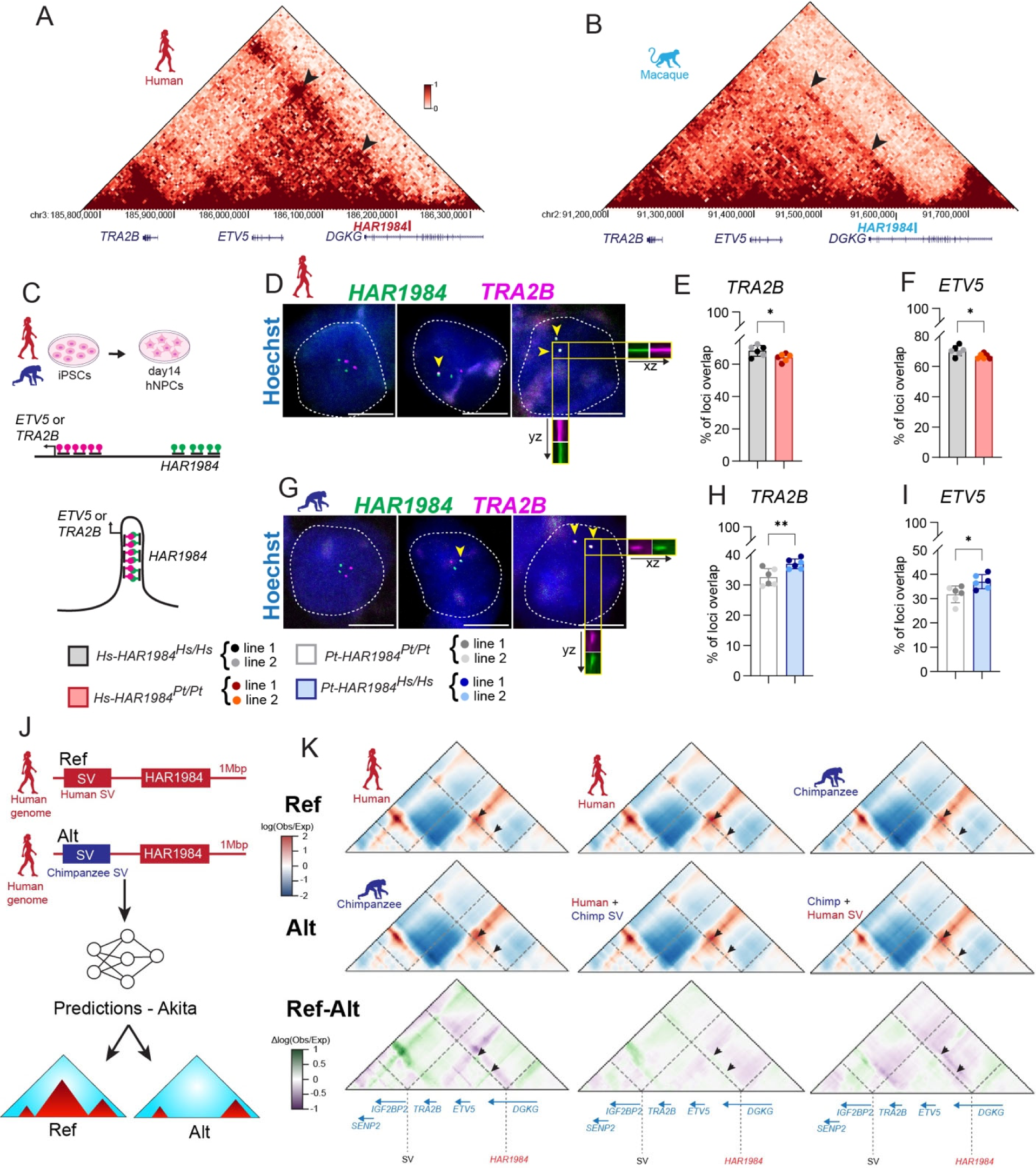
*Hs-HAR1984* locus promotes species-specific chromatin interactions with *ETV5* and *TRA2B*. (A, B) HiC map from human fetal brain (A) and macaque fetal brain (B) of *HAR1984* locus. Color scale represents normalized contact frequency. Arrowheads, *HAR1984-ETV5* or *TRA2B* contacts. (C) Experimental design of DNA FISH experiments. Human and chimpanzee control and *HAR1984* knock-in lines were differentiated to generate NPCs, with DNA FISH performed at D14. (D) Examples in human NPCs of no overlap between *HAR1984* (green) and *TRA2B* (magenta) loci (left), only one chromosome overlap (center), both chromosomes overlap (right). Example of xz and yz axis are shown of one overlapping chromosome. Yellow arrowpoints point to overlapping signal. (E-F) Quantification of *HAR1984-TRA2B* (*Hs-HAR1984^Hs/Hs^* n=112 chromosomes; *Hs-HAR1984^Pt/Pt^* n=132 chromosomes) (E) and *HAR1984-ETV5* (*Hs-HAR1984^Hs/Hs^* n=104 chromosomes; *Hs-HAR1984^Pt/Pt^* n=126 chromosomes) (F) overlap in human NPCs. (G) Examples in chimpanzee NPCs of FISH overlap between *HAR1984* (green) and *TRA2B* (magenta) loci, as in (D). (H-I) Quantification of *HAR1984-TRA2B* (*Pt-HAR1984^Pt/Pt^* n=144 chromosomes; *Pt-HAR1984^Hs/Hs^* n=138 chromosomes) (H) and *HAR1984-ETV5* (*Pt-HAR1984^Pt/Pt^* n=122 chromosomes; *Pt-HAR1984^Hs/Hs^* n=124 chromosomes) (I) overlap in chimpanzee NPCs. (J) Depiction of chromatin interaction predictions with SuPreMo-Akita 1Mbp region around *Hs-HAR1984* with human or chimpanzee SVs. (K) Hi-C maps of human or chimpanzee prediction with human SV365859. Ref (first row): Reference Hi-C data from human, human, and chimpanzee genomes. Color scale represents enrichment (red) versus depletion (blue) of chromatin interactions. Alt (second row): Hi-C chimpanzee, Predicted interactions for human genome with chimpanzee SVs, Predicted interactions for chimpanzee genome with human SVs. Differences between Ref-Alt (third row): Human vs chimpanzee genomes, human vs human with chimpanzee SVs, and chimpanzee vs chimpanzee with human SVs. Color scale represents chromatin interactions with predicted gain in Ref relative to Alt (green) and loss in Ref relative to Alt (purple). Arrows points to *HAR1984-ETV5* or *TRA2B* contacts. Scale bars: 10 μm. Each dot represents a single differentiation. N=3 differentiations (E-F, H-I). Unpaired t-test. \**p<*0.01; \*\**p<*0.001. Data are mean ± SD. See also Figure S7 and Table S4.

We next experimentally tested if *HAR1984* directly controls expression of its target genes across species via differential chromosome looping. For this we performed DNA FISH to test the extent to which human-specific variants in *HAR1984* modulate chromatin interactions between *HAR1984* and target gene promoters (Figure 6C). We used probes against *HAR1984* and the promoters of either *ETV5* or *TRA2B* in 2D differentiated human control and *Hs-HAR1984^Pt/Pt^* NPCs. Compared to human controls, *Hs-HAR1984^Pt/Pt^* NPCs exhibited reduced genomic interactions with both *ETV5* and *TRA2B* promoter regions (Figure 6D-F, S7A). The opposite was true in chimpanzee NPCs where the human sequence of *HAR1984* (*Pt-HAR1984^Hs/Hs^*) induced increased interactions between the enhancer and both target gene promoters, compared to the chimpanzee control (Figure 6G-I, S7B). These data demonstrate that the human sequence of *HAR1984* is both necessary and sufficient to contribute to stable chromatin interactions with *ETV5* and *TRA2B* promoters.

Species-specific larger genomic structural changes in regions adjacent to HARs have been shown to affect enhancer-promoter loops of these regions (Keough et al., 2023). To investigate how the genomic regions surrounding *Hs-HAR1984* might contribute to interactions with its target genes, we employed SuPreMo-Akita, a chromatin interaction prediction software (Fudenberg et al., 2020; Gjoni & Pollard, 2024). Akita is a convolution neural network (CNN) able to transform ∼1 megabase input DNA sequence into predicted chromatin interaction frequency maps (Figure 6J), and SuPreMo-Akita is a bioinformatics tool that predicts how genetic variants alter chromatin interactions. We identified 17 human-specific Structural Variations (SVs) in the region surrounding *Hs-HAR1984* (13 deletions and 4 insertions) (Table S4) (Kronenberg et al., 2018). Using SuPreMo-Akita, we modeled how chromatin interactions are impacted by the chimpanzee orthologue SV region computationally edited into the human genome and vice versa. We identified two human-specific deletions that favored chromatin contacts between *HAR1984* and *ETV5* and *TRA2B* promoters in humans compared to chimpanzees (Figure 6K and S7C). Presence of the chimpanzee SV is predicted to reduce these contacts (Figure 6K middle). These data suggest genomic co-evolution in which multiple human-specific sequence changes, including single nucleotide mutations within *HAR1984* and deletion of adjacent SV sequence, functionally converged to induce species-specific changes in chromatin conformation. Ultimately these genomic modifications help favor higher *ETV5* and *TRA2B* expression in the developing human cortex to promote neurogenesis.

## Discussion

The evolutionary molecular mechanisms by which uniquely human features are formed have been elusive. Our genomes contain thousands of human-specific loci, including HARs, with largely unknown functions. In this study we discover that *HAR1984* is an essential cis-regulatory element that shapes cortical development across species by modulating chromosomal interactions. By integrating *in silico, in vitro,* and *in vivo* approaches across human, chimpanzee and mouse models, we demonstrate that *Hs-HAR1984* functions as a transcriptional enhancer of two critical targets, *TRA2B* and *ETV5*. We discover a pivotal role of this enhancer in modulating chromosomal interactions to implement gene expression during human neurogenesis. By fine-tuning gene expression, *Hs-HAR1984* promotes neurogenesis leading to larger brains with more neurons and cortical folding (Figure 7). In sum, our study demonstrates essential roles for HARs in modulating chromosomal topology to influence human-specific development.

**Figure 7.**
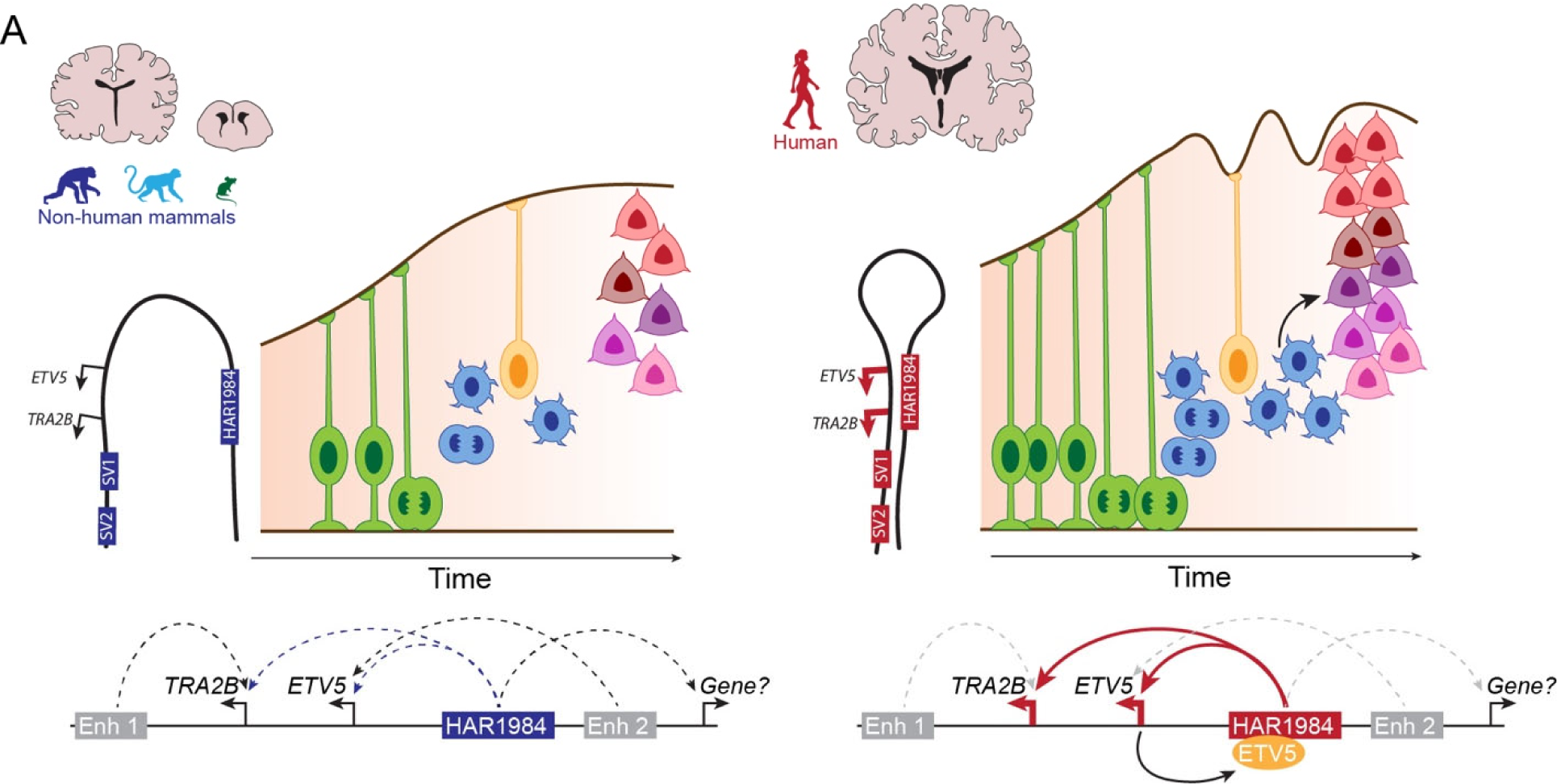
*Hs-HAR1984* promotes species-specific chromatin architecture and neurogenesis. (A) Summary of the study. *Hs-HAR1984* increases enhancer activity to promote *ETV5* and *TRA2B* expression amplifying NPC proliferation and neurogenesis, resulting in increased cortical size and folding. At the chromatin level, the frequency of interactions between *Hs-HAR1984* and promoters of *ETV5* and *TRA2B* are increased in humans versus non-human mammals. The model indicates *ETV5* and *TRA2B* may also have alternative enhancer regions (Enh1 and Enh2) that promote expression in non-human mammals, and with minor roles in humans. Moreover, ETV5 transcription factor binds *Hs-HAR1984* generating a positive feedback loop of regulation and leading to a robust activation of *HAR1984* direct and downstream targets and pathways.

We show that *Hs-HAR1984* contributes to cortical folding *in vivo,* illustrating that an enhancer can induce folding in the otherwise lissencephalic mouse cortex. This raises the question of how *Hs-HAR1984* influences cortical gyrification? Previous studies have reported increased folding in ferret, mouse, and marmoset models following genetic manipulations, including of human-specific genes (Dave et al., 2011; Heide et al., 2020; Hou et al., 2021; Matsumoto et al., 2020; Nonaka-Kinoshita et al., 2013; Singh et al., 2024; Stahl et al., 2013; Wang et al., 2016; Yabut et al., 2015). A common feature across these models is amplified basal progenitor number. Similarly, we show that *Hs-HAR1984* promotes TBR2+ basal progenitor number in humanized chimpanzee organoids and humanized mice. Reciprocally, we observe a dramatic reduction of TBR2+ cells in human organoids containing *Pt-HAR1984*. Taken together with previous studies, this reinforces a mechanistic link between basal progenitor amplification and gyrification. However, cortical folding may also result from interplay with additional factors such as cortical surface expansion, mechanical forces from the skull, and migratory dynamics (Del Toro et al., 2017; Javier-Torrent et al., 2021; Llinares-Benadero & Borrell, 2019; Walter et al., 2023). Indeed, *HAR1984* and its targets are also active and expressed in postmitotic neurons, raising the possibility that folding arises through both progenitor-mediated and neuron-intrinsic mechanisms.

*Hs-HAR1984* also contributes to cortical expansion, associated with amplified neuron production. This is mediated by *HAR1984* function in promoting neural progenitor proliferation and fate specification, with the most notable impact on basal progenitors. We further identify downstream pathways by which *HAR1984* may modulate neurogenesis. For example, we observe alterations in the Shh pathway, which may help explain shifts in neural cell fate and cortical folding in the *Hs-HAR1984* mouse model (Hou et al., 2021; Komada et al., 2008; Wang et al., 2016; Yabut et al., 2015). To date a number of human-specific coding and non-coding loci have been shown to modulate gene expression networks to promote cortical expansion (Fiddes et al., 2018; Fiddes et al., 2019; Fischer et al., 2022; Florio et al., 2018; Heide & Huttner, 2021; Liu et al., 2025; Pravata et al., 2025). Thus, multiple loci, including *HAR1984*, likely work in concert to contribute to human brain features including larger progenitor pools, increased neuronal output, and measurable increases in cortical size and folding.

We define *ETV5* and *TRA2B* as key direct regulatory targets of *HAR1984* during human neurogenesis. Our Hi-C and DNA-FISH experiments in human NPCs demonstrate that *Hs-HAR1984* mediates chromosomal interactions with *ETV5* and *TRA2B* loci to influence expression of both genes and their network of downstream targets. Moreover, independent depletion of *ETV5* and *TRA2B* phenocopies *HAR1984* perturbation of TBR2+ IPs. This is consistent with their central role downstream of *HAR1984* and well-established roles in neurogenesis (Fontanet et al., 2018; Roberts et al., 2014; Storbeck et al., 2014). Notably, both *ETV5* and *TRA2B* are expressed during neurodevelopment of non-human species (Di Bella et al., 2021; Zhu et al., 2018), suggesting that additional enhancers may contribute to their expression (Figure 7). We also discover that species-specific binding of ETV5 to *Hs-HAR1984*, enables a positive feedback loop to amplify enhancer activity. Such feedback regulation may underlie the robust and sustained effects of *HAR1984* on cortical development. Beyond ETV5, we identified additional species-specific transcription factors which may promote human-specific transactivation of *HAR1984*. (Table S4) The extent to which these trans factors are themselves targets of evolution is an interesting point for future study.

Our work demonstrates direct functions for a HAR in modulating chromatin topology and highlights how genomic structural variation can help facilitate genomic interactions. Notably, ∼30% of HARs fall within 500 kb of human-specific SVs that are predicted to reconfigure local chromatin loops and redirect enhancer–promoter interactions (Keough et al., 2023). Indeed, using chromatin interaction prediction models, we identified human-specific SVs adjacent to *Hs-HAR1984* that are predicted to enhance its looping interactions with target genes. New regulatory interactions may arise from modulation of TAD boundaries and CTCF binding sites, as evidenced by analyses of non-coding duplications and transposable element insertions (Choudhary et al., 2023; Franke et al., 2016). These human-specific sequences thus can promote enhancer– promoter looping, possibly by modulating trans factors. Together, this provides a compelling example of co-evolution between single nucleotide changes and SVs in regulatory evolution. This suggests potential mechanisms by which SVs act synergistically with HARs to modulate 3D genome topology and drive human-specific gene expression in cortical development.

Taken together, our results demonstrate that *HAR1984* is an essential transcriptional enhancer with both conserved and uniquely human roles in brain development. Its human-specific sequence boosts enhancer activity in neural progenitors by strengthening chromatin looping with neurodevelopmental genes, and establishes a feedback loop via ETV5 binding, leading to increased neurogenesis, which promotes cortical expansion and folding. These findings contribute to a growing body of work revealing how evolutionary changes in non-coding regulatory DNA underlie the emergence of human-specific traits.

## Material and Methods

### Animals

All mice experiments were approved by Duke IACUC and followed the guidelines from the Division of Laboratory Animal Resources from Duke University School of Medicine. Plug dates were defined as embryonic day (E) 0.5 on the morning the plug was identified. The following previously generated mouse lines were used in this study: *Tbr2*-2A-Flp knock-in and Flp-dependent Ai65-Frt reporter (Huilgol et al., 2023) with new mouse lines generated as described below. Genotyping primers specific to the mouse lines are listed in Table S5. No sample size calculation was performed. Data collection was performed by independent investigators, using double-blind and randomized analyses. Equivalent sexes were used (and not specified) and ages were indicated in study.

### Plasmid construction

The sequences of the PCR primers and synthetic oligonucleotides used for cloning are listed in Table S5. All sequences were validated by Sanger sequencing. All cloning PCR products were amplified by Q5 polymerase (NEB, M049L). For construction of vector used for *GFP-ETV5* overexpression, human ETV5 was amplified from human cortical organoid cDNA and inserted into the pCAGGS-EGFP vector by BsrGI enzyme site. For construction of vectors used for luciferase and IUE assays, human, chimpanzee HARs, and mouse *HAR1984* fragments were amplified from genomic DNA of individual species and inserted into the pNL1.1 vector (Promega) by KpnI and pPGK-EGFP (Addgene, #169744) by BglII. The fragments were inserted into the cut vector using HiFi DNA Assembly Master Mix (NEB, E2621L). For CRISPR-Cas9 knock-in of *Pt-HAR1984* in human ESCs and *Hs-HAR1984* in chimpanzee iPSCs, guide RNAs were designed using the online tool (Concordet & Haeussler, 2018) (http://crispor.tefor.net/). Two pairs of guide RNAs targeting both 5’ and 3’ side of *HAR1984* loci (*Hs-HAR1984* hg38 chr3:186,239,308-186,239,606; *Pt-HAR1984* panTro6 chr3:184,330,036-184,330,334), referred as guide1 and guide2. For cloning of the guides, briefly, the sense and antisense strand oligos for guide1 and guide2 were annealed and phosphorylated, and the duplexes were cloned into PX330 (Addgene, #158973) by BbsI site. Colonies were sequence validated using the U6_F primer. To increase the targeting efficiency, we amplified the whole sequence of U6_promoter up to the gRNA structure of guide1 and inserted it to guide2 vector to generate plasmid PX330-Guide1+2 by BamHI enzyme site. For construction of donor vectors used for CRISPR, human and chimpanzee *HAR1984* fragments were amplified from genomic DNA of individual species, left and right homology arms (∼800bp) were amplified from genomic DNA of individual species with PAM site mutated. PCR fragments were ligated by HiFi DNA Assembly Master Mix (NEB, E2621L) into the pCAG-Puro vector by XhoI enzyme site.

### Luciferase assay

HARs nomenclature is from (Girskis et al., 2021). Nomenclature used upon identification is listed in Table S1 ((Bird et al., 2007; Gittelman et al., 2015; Lindblad-Toh et al., 2011; Pollard et al., 2006; Prabhakar et al., 2006). Human NPCs differentiated from iPSC line (8799) (Miller et al., 2017) (Figure 1) or H9 ESC line (Figure 5), cells were split and plate at D10 by Accutase. 4×10^5^ cells were plating on 24-well plate coated with Poly-O-Ornithine/Laminin-coated. On day14, cells were transfected using Lipofectamine LTX reagent (Thermo Fisher Scientific, A12621) the following day with 250ng either mouse or chimpanzee and human mutation pNL1.1 luciferase vectors generated with enhancer sequences as described above. 250ng Firefly plasmid was co-transfected to control transfection efficiency. The luciferase assays were performed 24 hrs after transfection using the Nano-Glo Dual-Luciferase Reporter Assay System (Promega, N1551) according to the manufacturer’s instructions. Luciferase activity was measured and quantified by Luminescence Microplate Reader System. For each experiment, the raw value was normalized by pNL1.1 empty vector. Assay was performed from at least 3 independent differentiations with 3 technical replicated.

### *In utero* electroporation (IUE)

*In utero* electroporation was performed as previously described (Saito & Nakatsuji, 2001). Briefly, each E13.5 CD-1 embryo was injected with 1µl of plasmid mix (containing 0.01% fast green and 1,250ng of pPGK-HAR-EGFP and 750ng of pCAGGS-mCherry) and electroporation parameters: five 50 ms-pulses at 45V with 950 ms pulse-interval by platinum-plated BTX Tweezer Rodes. Plasmids were produced using Zymo Endotoxin Free Maxi Prep kits (Zymo, D4202) by following the manufacturers’ instructions.

### Transgenic LacZ mouse generation and β-gal activity assay

*Hs-HAR1984::LacZ* mice were generated at the Duke Transgenic Mouse facility using standard gene targeting techniques on B6SJLF1/J hybrid mouse ES cells. *Hs-HAR1984* was cloned in Hsp68-LacZ (Addgene, #37843) by XhoI-Ascl enxyme sites. DNA for pronuclear injection was prepared by digesting fully constructs with XhoI/MfeI (NEB) and purified by gel extraction (Zymoclean, 11-300C). All linearized constructs were submitted to the Duke Transgenic Mouse Facility for pronuclear injection into B6SJLF1/J strain blastocysts (F1 hybrid from C57BL/6J X SJL/J). Transgenics were genotyped (primers Table S5) and two independent founders were backcrossed to C57BL/6J and used for subsequent analysis and breeding. To test for enhancer activity, we used a standard enhancer assay for β-Galactosidase activity. In brief, E12.5, E14.5 or E16.5 embryos were dissected in PBS and place in ice-cold fix solution (0.02% IGEPAL, 1% formaldehyde, 0.2% glutaraldehyde in PBS) for 2hrs with shaking. Embryos were permeabilized with three washed in wash solution (2mM MgCl_2_, 0.02% IGEPAL in PBS) at room temperature. Staining was performed overnight in β-gal solution (12mM K-Ferricyanide K3Fe(CN)_6_, 12mM K-Ferrocyanide K4Fe(CN)_6_, 0.02% IGEPAL, 4mM MgCl_2_, 16ug of X-gal (Thermo Fisher Scientific, 15520034) in PBS) on shaker in 37°C incubator. After staining, two 30min washed were performed in wash solution. Embryos were stored in storage buffer (4% formaldehyde, 100mM Sodium Diphosphate, 10% methanol in PBS) at 4°C. Images of whole-mount LacZ were obtained on a Leica M165 FC microscope using the Leica Application Suite software package (v.4.1.0). Two independent *Hs-HAR1984::LacZ* lines were analyzed.

### Cell line maintenance and NPC differentiations

All lines tested negative for mycoplasma contamination in this study were checked by the MycoAlert Mycoplasma Detection Kit (Lonza, LT07-518). One human ESC line WA09 (H9), one human iPSC line (8799), one chimpanzee iPSC line (C3649) were used in this study. H9 (WA09) was purchased from WiCell (WB68167). 8799 was used in this study was previously characterized and genotyped (Miller et al., 2017). C3649 was a gift from Yoav Gilad (Gallego Romero et al., 2015). ESCs and iPSC lines were supplemented with mTesR^TM1^ (Stem Cell, 85850) with Normocin (Invitrogen, ant-nr-1). All cells were cultured at 37°C in 5% CO2 environment.

All ESCs (including control and CRISPR KI) and iPSCs (including 8799, C3649 and CRSIPR KI) were cultured in Matrigel-coated plates with mTeSR^1TM^ media (Stem Cell, 85850) in an undifferentiated state. Cells were passaged at a 1:5 ratio by treatment with ReLeSR (Stem Cell, 5872)/Accutase (Thermo, A11105) for maintaining.

For luciferase assay experiment, 8799 iPSCs or H9 (ESC) were differentiated to neural progenitor cells (NPCs) as previously described (Shi et al., 2012). Briefly, 0.8×10^6^ cells were seeded on Matrigel coated 60mm plates until reaching 100% confluent (Day 0), medium was replaced with neural induction medium containing 1μM Dorsomorphin (Sigma, P5499), 10μM SB431542 (Selleckchem, S1067) every day. On Day 6, cells were washed with DMEMF12, dissociated with 1mg/mL DispaseII (Thermo, 17105041), clumps suspended in neural maintenance medium and plated on Poly-O-Ornithine (Sigma, P4957) /Laminin (Sigma, L2020) coated 60mm dishes. After D8 when neural rosettes appeared, 20ng/mL bFGF2 (R&D,233FB025) was added in neural maintenance medium.

For chimpanzee NPCs differentiation, C3649, *Pt-HAR1984^Pt/Pt^* control, *Pt-HAR1984^Hs/Hs^ −1* and *Pt-HAR1984^Hs/Hs^ −2* cells were differentiated to neural progenitor cells (NPCs) as described (Keough et al., 2023; Whalen et al., 2023). Briefly, 0.7×10^5^ cells were seeded on Matrigel coated 12-well plate (Day 0), medium was replaced with N2-B27 neural induction medium containing 50 ng/mL bFGF2(R&D,233FB025), 100 ng/mL Noggin (Miltenyi Biotec,130-103-456) every day until collection.

### Human and chimpanzee CRISPR mutagenesis

For establishment of the *Hs-HAR1984^Pt/Pt^* mutant lines, plasmids PX330-hsguide1+2 (1µg) and pCAG-Puro-Pt-HAR1984 (2µg) were electroporated into 1×10^6^ H9 cells using the P3 Primary Cell 4D-Nucleofector kit (Lonza, V4XP-3024). Following each electroporation, cells were separated to grow in 3 wells of a Matrigel (VWR, 354277) coated 6-well plate in mTesR^TM1^ supplemented with 1x CloneR (Stem Cell, 5888). After 3 days from electroporation, the cells were selected with 0.5 mg/ml Puromycin (Sigma, P8833). After approximately 5 days in selection medium, cells were switched back to mTesR^TM1^ medium until the single colony was big enough to passage. Single colonies were picked to Matrigel coated 24-well plates. After about 6 days, when the colonies reach ∼70% confluent in the well, colonies were collected half for freezing and half for genotyping after dissociated by ReLeSR (Stem Cell, 5872). For the colonies genotyping, genomic DNA were extracted by Genomic DNA Purification Kit (Stem Cell, 79020) and Sanger sequencing was used to screen for correct gene editing. Primers listed in Table S5. A total of 2 homozygote *Hs-HAR1984^Pt/Pt^* lines were generated from ∼150 colonies. Homozygote colonies were tested Karyotypes through hPSC Genetic Analysis Kit (Stem Cell, 7550).

For establishment of the *Pt-HAR1984^Hs/Hs^* lines, plasmids pX330-ptGuide1+2 (1µg) and pCAG-Puro-Hs-HA1984 (2µg) were electroporated into 1×10^6^ C3649 cells using the P3 Primary Cell 4D-Nucleofector kit (Lonza, V4XP-3024). The selection and genotyping process was the same as above. A total of 2 homozygote *Pt-HAR1984^Hs/Hs^* lines were generated from ∼150 colonies. Pluripotency was tested through *OCT4* RT-qPCR. Primers listed in Table S5.

### Cortical organoids generation and dissociation

Forebrain human cortical organoids were generated as described (Velasco et al., 2019). Briefly, on day 0, H9, *Hs-HAR1984^Hs/Hs^* control, *Hs-HAR1984^Pt/Pt^-1*, *Hs-HAR1984^Pt/Pt^-2*, or BFP-dCas9-KRAB cells were dissociated to single cells by Accutase, with 9,000 cells per well were reaggregated in ultra-low cell-adhesion 96-well plates with V-bottomed conical wells (S-bio, MS-9096VZ) in Cortical Differentiation Medium (CDM) I (20% KSR, 0.01mM NEAA, 1mM pyruvate, 0.1mM β-mercaptoethanol, Pen/strep in Glasgow MEM). Day 0-6, ROCK inhibitor Y-27632 (Stem Cell, 72304) was added (final concentration of 20 µM). Day 0-18, Wnt inhibitor IWR1 (Millipore, 681669) and SB43154 (Selleckchem, S1067) were added (final concentration of 3 and 5 µM, respectively). From day 18, the aggregates were cultured in ultra-low attachment culture dishes under orbital agitation in CDM II (1x Glutamax, 1x N2 supplement, 1% chemically concentrate lipids, Pen/strep in DMEM/F12) until day 35 and in CDM III (1x Glutamax, 10% FBS, 1x N2 supplement, 1% chemically concentrate lipids, 0.5 µg/ml heparin, Pen/strep in DMEM/F12) until day 70 and, and CDM IV (1x Glutamax, 10% FBS, 1x N2 supplement, 1x B27 supplement without vitamin A, 1% chemically concentrate lipids, 0.5 µg/ml heparin, Pen/strep in DMEM/F12) until day 90.

Forebrain chimpanzee organoids were generated as described above for human cortical organoids with few changes. Briefly, on day 0, C3649, *Pt-HAR1984^Pt/Pt^* control, *Pt-HAR1984^Hs/Hs^-1*, *Pt-HAR1984^Hs/Hs^-2* cells were dissociated and seeded in the same plates and same confluency described for the human cells in Cortical Differentiation Medium (CDM) I. Day 0-6, ROCK inhibitor Y-27632 (Stem Cell, 72304) was added (final concentration of 20 µM). Day 0-18, Dorsomorphin (Sigma, P5499) and SB43154 (Selleckchem, S1067) were added (final concentration of 10 and 5 µM, respectively). From day 18, the chimpanzee aggregates were growth as described above for the human aggregates.

Dissociation of organoids in 2D cultures were performed as described in (Miura et al., 2020) with minor modifications described as follow. BFP-dCas9-KRAB was dissociated at D55. Thus, 5 x 10^4^ cells were seeded in wells of 24-well glass bottom plates (Mattek, P24G-1.5-10-F) coated with poly-D-lysine (Sigma-Aldrich, P7280-5MG). Lentivirus delivering sgRNAs were added 3-4hrs after dissociation to allow cells to attach. Cultures were maintained in CDM III for 5 days, then fixed for 15 min at room temperature with 4% paraformaldehyde (PFA) in PBS or collected for RNA extraction.

### Immunofluorescence staining

For histological analysis, samples (including mouse embryos, organoids, and NPCs) were collected and dissected in cold phosphate buffer saline (PBS), postnatal 30 and adult animals were performed with trans cardiac perfusion by PBS before dissection. Brains were fixed by immersion in a 4% Paraformaldehyde/PBS solution, the fixation time varied due to samples (overnight for embryonic brains at 4°C, 4hrs for adult brains at 4°C, 3hrs for organoids at 4 °C, 15mins for NPCs at room temperature). Following fixation, embryonic and adult brains were cryoprotected overnight in 30% sucrose/PBS solution. Coronal cryosections were collected using a cryostat with a variable thickness depending on experiments (20mµ for of coronal sections of embryo brains, 30µm for sagittal sections of embryonic brains and adult coronal sections). Immunofluorescence was performed as described previously. Briefly, slides were permeabilized with a 10min wash in 0.3 % Triton-X/PBS. After blocking sections in 5% NGS/PBS for 1hr at room temperature, primary antibody incubation in PBS was performed overnight at 4°C and secondary antibody for primary incubation for 1hr at room temperature. Antibody staining was listed below: SOX2 (Thermo,14-9811-82,1:500); TBR2 (Abcam, ab183991,1/500, for mouse sections); TBR2 (Abcam, ab23345, 1/300, for organoids sections and human NPCs); PPP1R17 (Sigma-Aldrich, HPA047819, 1/150); CTIP2 (Abcam, ab18465,1/500); TBR1 (Abcam, ab31940,1/500); PH3 (Cell Signaling, 9706,1/1000); CUX1 (Abcam, ab54583, 1/100, required citrate boiled 15min); LHX2 (Millipore, abe1402, 1/500); KI67(Cell signaling, 9449, 1/300); B-gal (Promega, Z3781, 1/50, required citrate boil 15min) RFP (Rockland, 600-401-379,1/2000); NEUN (Cell Signaling, 24307,1:300).

### Image acquisition and quantification

Images were captured using a Zeiss Axio Observer Z.1 equipped with an Apotome for optical sectioning at 10X, 20X with 4μm Z-stack. P30 RFP images were captured from Zeiss710 confocal for optical sectioning at 20X or 5X with no Z-stack. DNA FISH images were captured from Zeiss780 confocal for optical sectioning at 100X with 3µm Z-stack. Images were captured with identical exposures between genotypes. Some images were captured using tiling with higher magnification objectives to have higher resolution which in some cases creates appearance of artificial lines.

For each experiment, 2–4 sections were imaged per embryo or per organoid; brain section images were cropped (100μm radial columns for embryonic brains, 300μm radial columns for adults), and brightness adjusted equivalently across all images in Fiji. Cell counting of embryonic brain sections, human and chimpanzee organoids and NPC was performed in FIJI (ImageJ version Java1.8.0), using the Cell counter plugin (Schindelin et al., 2012). For organoids, we used 20x objectives with tiling to image the whole organoid, then quantified the whole field. Hoechst quantification was done by QuPath automatically cell counter (Bankhead et al., 2017) (version-0.4.2 and 0.5.1) automatically cell count combined with ImageJ. QuPath quantification parameters were adjusted as follows: requested pixel size = 0.1 µm, background radius = 5 µm, minimum area = 12 µm^2^, maximum area = 200 µm^2^, cell expansion = 3 µm, include cell nucleus and smoother boundaries boxes were unchecked. The threshold was set independently for each channel and maintained for each image in an experiment. Raw quantification data were initially processed by Excel (Microsoft version 2501) and then followed analyzed and graphed with Prism (GraphPad Software10).

### RNA extraction and RT-qPCR

Mouse embryonic cortices were dissected in cold PBS and RNA was purified from Trizol (Thermo, 15596026) method. For NPCs and organoids, RNA was purified using RNeasy micro-plus kit (Qiagen,74034). For mouse sample, 5ng cDNA was used for each qPCR reaction. For NPC and organoids characterizing, 15ng cDNA was used for each qPCR reaction. cDNA was prepared using iScript kit (Bio-Rad, 1708891) following manufacturer’s protocol. qPCR was performed using SYBR Green iTaq (Bio-Rad, 1725124). In at least three independent biological samples in a QuantStudio 6 machine (Applied System). Primers were listed in Table S5.

### Single-cell RNA sequencing and analysis

2-3 D60 or D90 human forebrain organoids from the following genotypes H9, *Hs-HAR1984^Hs/Hs^* control, *Hs-HAR1984^Pt/Pt^-1* and *Hs-HAR1984^Pt/Pt^-2* were dissociated into a single-cell suspension using the Worthington Papain Dissociation System kit (Worthington Biochemical, LK00315) as previously (Velasco et al., 2019). In brief, organoids were transfer in papain/DNase solution and minced in small pieces using a razor blade. The plates were incubated at 37°C on orbital shaker for 30min. Organoids were dissociated again in 1mL pipette and returned at 37°C for ulterior 10min. Papain inhibitor were then added to stop enzymatic dissociation. Cells were then centrifuged at 300g for 7min and then filtered (Sysmex celltrix, 04-004-2326). Dissociated cells were resuspended in ice-cold PBS containing 0.04% BSA (VWR, 97061-420) at a concentration of 1,000 cells/μl, and approximately 16,000 cells per channel (to give estimated recovery of 10,000 cells per channel) were loaded onto a Chromium Single Cell 3′ Chip (10x Genomics, PN-120236) and processed through the Chromium controller to generate single-cell gel beads in emulsion (GEMs). scRNA-seq libraries were prepared with the Chromium Single Cell 3′ Library & Gel Bead Kit v.2 (10x Genomics, PN-120237). Libraries from different samples were pooled based on molar concentrations and sequenced on a NovaSeq 6000 instrument (Illumina).

The 10x Genomics Cell Ranger Count command version 6.1.2 was used to align reads from RNA-seq to the GRCh38 human reference genome and generate filtered count matrices. The count matrices were further processed using the Seurat R package version 4.3.3 (Gokhman et al., 2021). Cells expressing a minimum of 500 genes and less than 5% mitochondrial genes were kept, and UMI counts were normalized for each cell by the total expression, multiplied by 10^6^, and log-transformed. Seurat’s default method for identifying variable genes was used with x.low.cutoff = 1, and the ScaleData function was used to regress out variation due to differences in total UMIs per cell. Principal component analysis (PCA) was performed on the scaled data for the variable genes, and top principal components were chosen. The datasets were integrated using the IntegrateData function. UMAP was generated with the RunUMAP function based on the top 50 principal components. Clusters were obtained using the integrated counts with the FindNeighbors function, based on the 50 principal components. The function FindClusters was used with resolution=1.1. Marker genes were obtained using the FindConservedMarkers function. Clusters cell identity was assigned according to previously used markers (Uzquiano et al., 2022) (Cycling cells: cluster 17, 18, 28, 29; aRGCs: clusters 16, 19, 20, 23, 25, 26, 31, 32, 34, 35, 36; oRGCs: clusters 2, 24, 33; IPs: clusters 11, 15; Immature neurons: clusters 3, 6, 8, 12, 13, 14, 22, 27, 30; Deep layers: clusters 1, 4, 7,9, 10; Upper layers: clusters 0, 5, 21). Cluster composition was calculated on total number of cells per cluster per sample. To obtain cluster-specific differentially expressed genes between *Hs-HAR1984^Pt/Pt^* and control, a pseudo-bulk approach was used (Lun et al., 2016; Robinson et al., 2010). The pseudo BulkDGE function from the scran R package version 1.22.1, which performs the edgeR approach for each cluster, was used. An FDR threshold of 0.05 with Log2(FC) ≥ 1.5 or ≤-1.5 was used to obtain differentially expressed genes (Table S2) (Charrier et al., 2012).

Pseudotemporal ordering of cells was performed using Monocle3 (v1.4.26) (Qiu et al., 2017) to infer lineage trajectories from scRNA-seq data. Seurat objects were first subset by genotype and converted into Monocle’s CellDataSet format using the raw count matrix, cell metadata, and gene annotations. Preprocessing was performed with preprocess_cds using 50 dimensions. To preserve prior dimensionality reduction, UMAP embeddings from the Seurat analysis were transferred directly into Monocle and used for downstream steps. Cells were clustered with cluster_cells, and the principal trajectory graph was learned with learn_graph. Cells were ordered in pseudotime using manually defined root nodes corresponding to early progenitor states based on UMAP visualization. Pseudotime distributions across samples were visualized as stacked bar plots after binning cells along the trajectory.

To evaluate and control for cell cycle–related transcriptional variation, we performed cell cycle scoring and regression using the Seurat package (v4.3.3) based on canonical S and G2/M phase marker genes from (Tirosh et al., 2016). Cell cycle phase scores were assigned using the CellCycleScoring function. PCA was performed on cell cycle genes to visualize phase separation, and cells were stratified into G1, S, or G2/M phases. Phase-specific marker expression (e.g., *PCNA*, *TOP2A*, *MCM6*, *MKI67*) was examined using ridge plots, and relative proportions of cells in each phase were quantified per sample. These distributions were visualized as stacked bar plots to assess sample-specific differences in cell cycle dynamics.

### Lentivirus packaging and transduction

HEK293T cultured in DMEM+10%FBS+1%P/S were used for lentivirus production. 26.75ug pKLO5-sgRNA-EGFP or -mScarlet (Addgene, #57822), 20ug Packaging plasmid (psPAX2, Addgene, #12260) and 6.25ug Envelope plasmid (pMD2.G, Addgene, 12259) were transfect with PEI-MAX (Polysciences, 24765-100) when cells reached to 70-80% in 15cm dish. 48hrs after transfection, the medium was collected and centrifuged at 3000g x 10min. The supernatant was then filtered through a 0.45mm filter (VWR, 28143-352) into an ultracentrifuge tube (Beckman-Coulter, 357448). 4ml sterile 20% sucrose was added below the medium using a 5 ml serological pipette. The mixture was centrifuge at 19700 rpm for 2hrs at 4°C in a swinging bucket rotor (Beckman-Coulter, SW28). The supernatant was then removed, 100µl of PBS was added, and the virus was resuspended overnight at 4°C with rocking. The resuspended virus was aliquoted, flash-freeze and stored at −80°C.

### Bulk RNA sequencing and analysis

For the mouse brain samples, E16.5 *Mm-HAR1984^Mm/Mm^*, *Mm-HAR1984^Hs/Mm^* and *Mm-HAR1984^del/del^* cortices were dissected in cold PBS and RNA was purified using RNeasy plus kit (Qiagen, 74034). For each genotype 3 biological replicates from two litters were sequenced. cDNA libraries were prepared by Illumina TruSeq stranded mRNA kit. RNAseq libraries were sequenced on the NextSeq2000 (PE100) with 20M paired end reads, to a depth of ∼40 million total reads per sample. Reads were aligned to the mm39 genome using STAR (Dobin et al., 2013), Differential gene expression (DGE) analysis using DESeq2 (Best et al., 2014), comparing genotypes. DGE lists were defined using an FDR <0.05, with Log2(FC) ≥ 1.3 or ≤-1.3 (Table S3). Overlapped and specific gene expression changes were defined by Biovenn (https://www.biovenn.nl/). Plot were made by R Script ggplot.

### iGONAD

*Mm-HAR1984^Hs/Mm^* and *Mm-HAR1984^del/del^* mouse lines were generated with iGONAD as previously described in (Gurumurthy et al., 2019). In brief, CRISPR mix components (6.1µM CAS9 (IDT, 1081058), 30µM gRNA (Alt-R® CRISPR-Cas9 sgRNA)) and 20µM ssDNA repair templates in Opti-MEM (Gibco, 31985070) were incubated at 37°C for 10min. Around 0.5µl of CRISPR mix components containing 0.01% fast green were injected into the ampulla of the oviducts of pregnant dams at E0.7. The oviducts were then electroporated to deliver the components to the developing zygote. The electroporator used is the CUY21EDIT II (BEX Co., LTD.) with these settings for C57BL/6J mice: Square mA mode, Pd V: 60 V, Pd A: 100 mA, Pd on: 5 msec, Pd off: 50 msec, Pd N: 3, Decay: 10%. See table S5 for sgRNAs sequences. All obtained pups were then screened with Sanger sequencing for successful editing. For the synthesis of ssDNA donor, the following coordinates were used: homology arm 1: mm39 chr16:22363786-22363985; homology arm 2: mm39 chr16:22364164-22364363; Hs-HAR1984: hg38 chr3:186239308-186239606.

### Cell cycle exit

For human organoids cell cycle exit, EdU (Thermo, A10044) was dissolved in PBS for 10mg/mL stock. At D58, single organoids were moved to 24-well ultra-low attachment plate and 20µM EdU were added in the CDM II. Organoids were harvested exactly 48 hr later at D60. EdU was detected by Click-iT® Plus EdU Imaging Kits (Thermo, C10638). Antibody staining was then performed as described above.

### Transcription factor prediction

Prediction for putative transcription factor binding sites on *Hs-HAR1984* or *Pt-HAR1984* was performed using the software QBic (Quantitative Predictions of TF Binding Changes due to sequence Variants) (Martin et al., 2019). The software uses ordinary least square (OLS)-based prediction model to assess impact of non-coding mutations of TF-DNA interactions (Zhao et al., 2017). The model is trained on high-throughput from multiple species. All the available datasets were used for the predictions to take into consideration species trans environment (see Table S4). Human sequence of *HAR1984* was use as input (hg38 chr3:186239308-186239606) and coordinates with single chimpanzee mutations were used as variants (hg38 chr3: 186239338 C>A; chr3:186239358 G>A; chr3: 186239374 G>A; chr3: 186239418 G>C; chr3: 186239429 T>C).

### Chromatin interaction prediction

Human specific SVs within 900 bp of HAR1984 were evaluated using SuPreMo-Akita. In short, a 1 Mb sequence centering the SV-HAR pair was inputted into the Akita model which outputs normalized pairwise contacts between all 2,048 bp bins in the input sequence. For each SV, three pairs of sequences were compared: hg38 vs. panTro6, hg38 vs. hg38 with chimpanzee deletion, and panTro6 vs. panTro6 with human insertion. The resulting map pairs were subtracted and the effect of each SV was inspected across the visualize the effect of the SV on genomic contacts.

### DNA FISH

Day 14 human (H9, *Hs-HAR1984^Hs/Hs^* control, *Hs-HAR1984^Pt/Pt^-1*, *Hs-HAR1984^Pt/Pt^-2)* and chimpanzee (C3649, *Pt-HAR1984^Pt/Pt^* control, *Pt-HAR1984^Hs/Hs^-1*, *Pt-HAR1984^Hs/Hs^-2*) NPCs were probed as previously described (Bayani & Squire, 2004) with some changes. Briefly, slides fixated in ice-cold MeOH for 20min at −20°C, then were treated with 10% pepsin (Abcam, ab64201) in pre-warmed 10mM HCl for 7min at room temperature. After a wash in PBS, the samples were passed through serial dehydrating series of 70%, 80% and 100% EtOH for 5min each. The slides were air-dried for another 5min. Samples were denatured in 10% formamide/2xSSC (pH 7.0) at 72°C for 2min then promptly places in ice-cold 70% EtOH for dehydration series with 80% and 100% EtOH for 5min each. Slides were then air-dry and proceed with probes hybridization. For probes were prepared and labeled with ClickTech DNA FISH Kit (Baseclick, BCK-DNA-FISH-488 and BCK-DNA-FISH-555) following manufacturer instructions. List of primers can be found in Table S5. Probes were diluted to a final concentration of 2ng/µL in hybridization buffer (62.5% Formamide, 2.5x SSC, 12% dextran Sulfate). Labeled probes were denatured for 5minutes at 75°C and immediately quick-chill on ice to avoid re-annealing. Total of 60ng of each probe was applied to each sample and then covered with hybridization cover slip (Silgma-Aldrich, GBL716024) and edges were closed with parafilm to avoid evaporation. Slides were incubated in humid chamber for 72hrs at 37°C in hybridization over. Following washes were performed at 45°C: 3x 5min in 50%formamide/2xSSC; 3x 5min in 1xSSC; 3x 5min 0.1% tween/4xSSC. Slides were mounted with Vectashield (VWR, 101098-042). For all samples and differentiations, only cells were all 4 loci (*HAR1984* and *ETV5* or *TRA2B*) were present were considered for quantifications.

### Statistics and reproducibility

Mouse samples were selected unbiased for various experimental conditions, samples were excluded from severely damaged or dead, or if the immunostaining signal or IUE efficiency was clearly lower than normal or was not comparable between the different groups of samples. Mouse experiment was performed from at least 2 independent litters with at least 3 animals for each condition, the sample sizes, biological and duplications are listed in the figure legends. Images shown in all figures are representative of at least 3 independent brains.

Human/chimpanzee cortical organoids samples were selected based on the RT-qPCR of *FOXG1/PAX6/OCT4* levels, samples were excluded based on the extremely not comparable *FOXG1* level from each differentiation. We only included organoids in which *FOXG1* levels were 800-fold higher than day 0 ESCs (almost all organoids fit this threshold). All analyses were performed blindly by 1 or more investigators. The sample sizes are listed in the figure legends. The number of data points, biological replicates and statistical tests used for all the comparisons are indicated in the figure legends. Human and chimpanzee cells experiments were performed from at least 3 independent differentiations, the sample sizes, biological and duplications are listed in the figure legends. Images shown in all figures are representative of at least 3 independent experiments.

The parametric tests used were two-tailed unpaired or paired Student’s t-test; one-way analysis of variance (ANOVA) followed by Dunnett’s multiple comparisons test; or two-way ANOVA followed by Bonferroni’s post-hoc test; Wilcoxon rank-sum test. The statistical test used for each experiment was specified in the figure legends. The level of statistical significance, statistical tests and sample size (n) for each experiment are informed in figures legends. Graphs and bar plot, means ± S.D. *p<0.01, **p<0.01, ***p<0.001, ****p<0.0001

## Supplemental information

**Figure S1.**
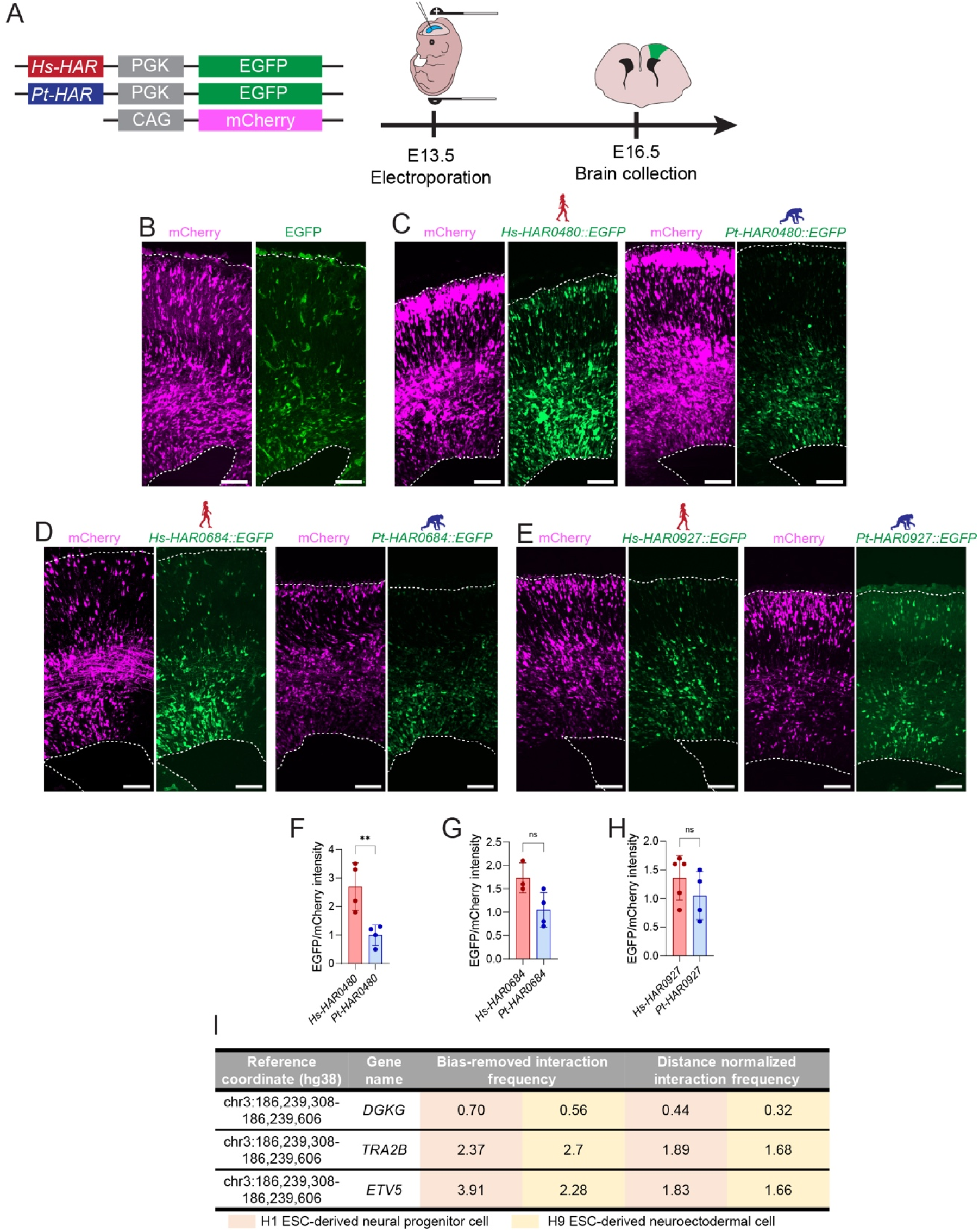
HAR candidates activity in vivo, related to Figure 1. (A) Depiction of plasmid used for *in utero* electroporation (left) and experimental design (right). pCAGGS-mCherry and *Hs-HAR::EGFP* or *Pt-HAR::EGFP* were injected in the ventricles of embryos at E13.5. Brains were collected and analyzed at E16.5. (B-E) Representative images of control (only promoter) (B), and enhancer candidates (C-E). (F-H) Quantifications of intensity of EGFP relative to mCherry for C-E. (I) HiC data showing *ETV5* and *TRA2B* is the highest frequency target of *HAR1984* in human neural cells. Each dot represents single embryos from 2 independent litters. Unpaired t-test (F-H). ns, not significant; \*\**p<*0.001. Data are mean ± SD.

**Figure S2.**
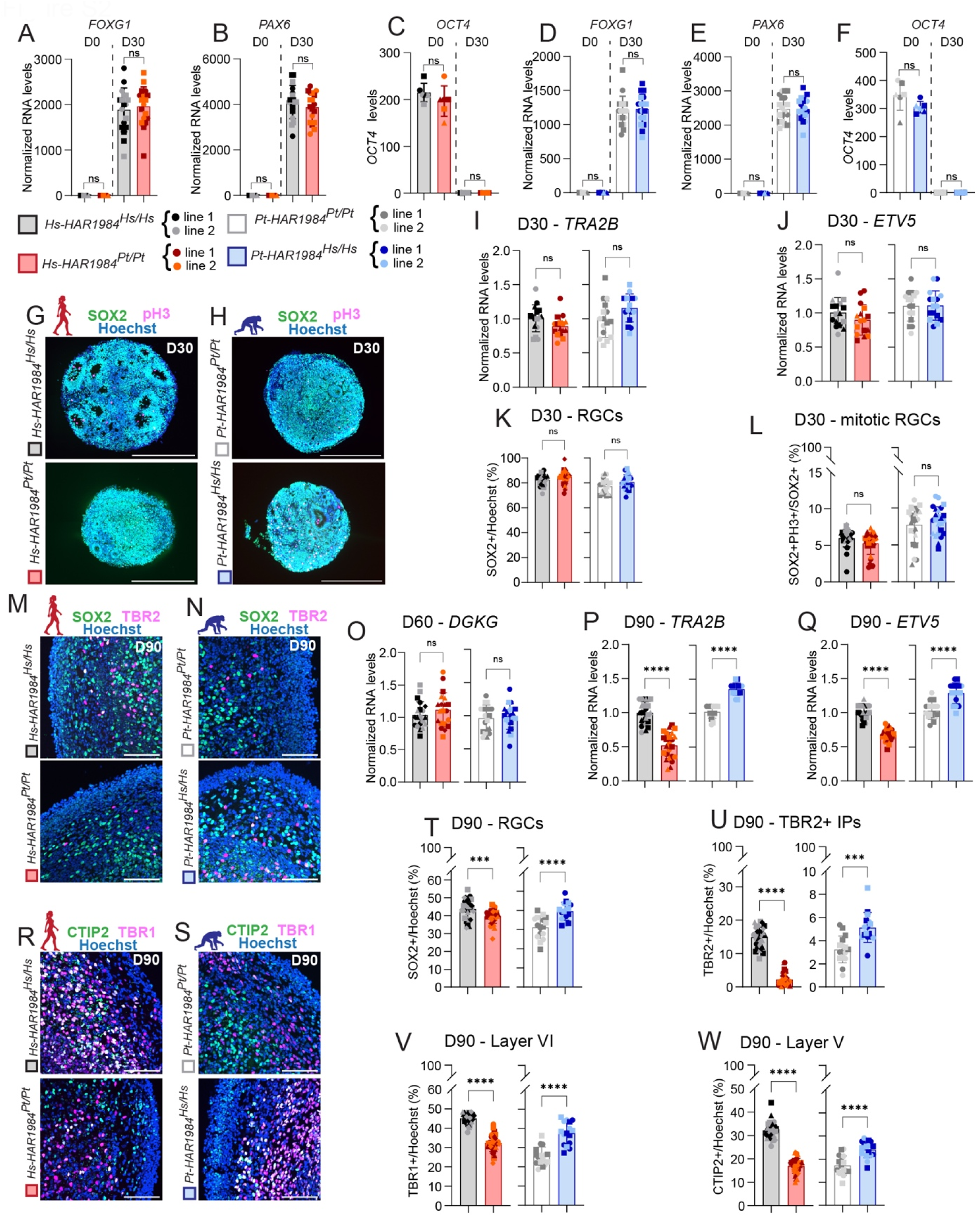
*Hs-HAR1984* promotes neurogenesis at late, but not early stages of development in human and chimpanzee cortical organoids, related to Figure 2. (A-C) RT-qPCRs for cortical markers *FOXG1* (A) and *PAX6* (B), and pluripotency marker *OCT4* (C) and D0 and D30 in human cortical organoids. (D-F) RT-qPCRs of cortical markers *FOXG1* (D) and *PAX6* (E), and pluripotency marker *OCT4* (F) and D0 and D30 in chimpanzee cortical organoids. (G-H) Immunostaining for SOX2 (RGCs, green) and PH3 (mitotic cells, magenta) in D30 human (G) and chimpanzee (H) organoids. (I-J) RT-qPCR of *TRA2B* (I) and *ETV5* (J) levels normalized on control in human and chimpanzee D30 organoids. (K) Quantification of SOX2 percentage over total number of cells (Hoechst) shown in G-H. (L) Quantification of SOX2+PH3+ percentage over total number of RGCs (SOX2) shown in G-H. (M-N) Immunostaining for SOX2 (RGCs, green) and TBR2 (IPs, magenta) in D90 human (M) and chimpanzee (N) organoids. (O) RT-qPCR of *DGKG* levels normalized on control in human and chimpanzee D60 organoids. (P-Q) RT-qPCR of *TRA2B* (P) and *ETV5* (Q) levels normalized on control in human and chimpanzee D90 organoids. (R-S) Immunostaining for CTIP2 (layer V neurons, green) and TBR1 (layer VI neurons, magenta) in D90 human (R) and chimpanzee (S) organoids. (T-W) Quantification of SOX2 (T), TBR2 (U), TBR1 (V) and CTIP2 (W) percentage over total number of cells (Hoechst) shown in M-N and R-S. Scale bars: 500 μm (G-H) 100 μm (M-N, R-S). Each dot represents a single organoid. Different dot shapes represent different batches of differentiation. D30 and D60 human organoids: 4 differentiations. D30 chimpanzee organoids and D90 human organoids: 3 differentiations. D90 chimpanzee organoids: 2 differentiations. 3-4 organoids were analyzed in each differentiation. Unpaired t-test. ns, not significant; \*\*\**p<*0.0001; \*\*\*\**p<*0.0001. Data are mean ± SD.

**Figure S3.**
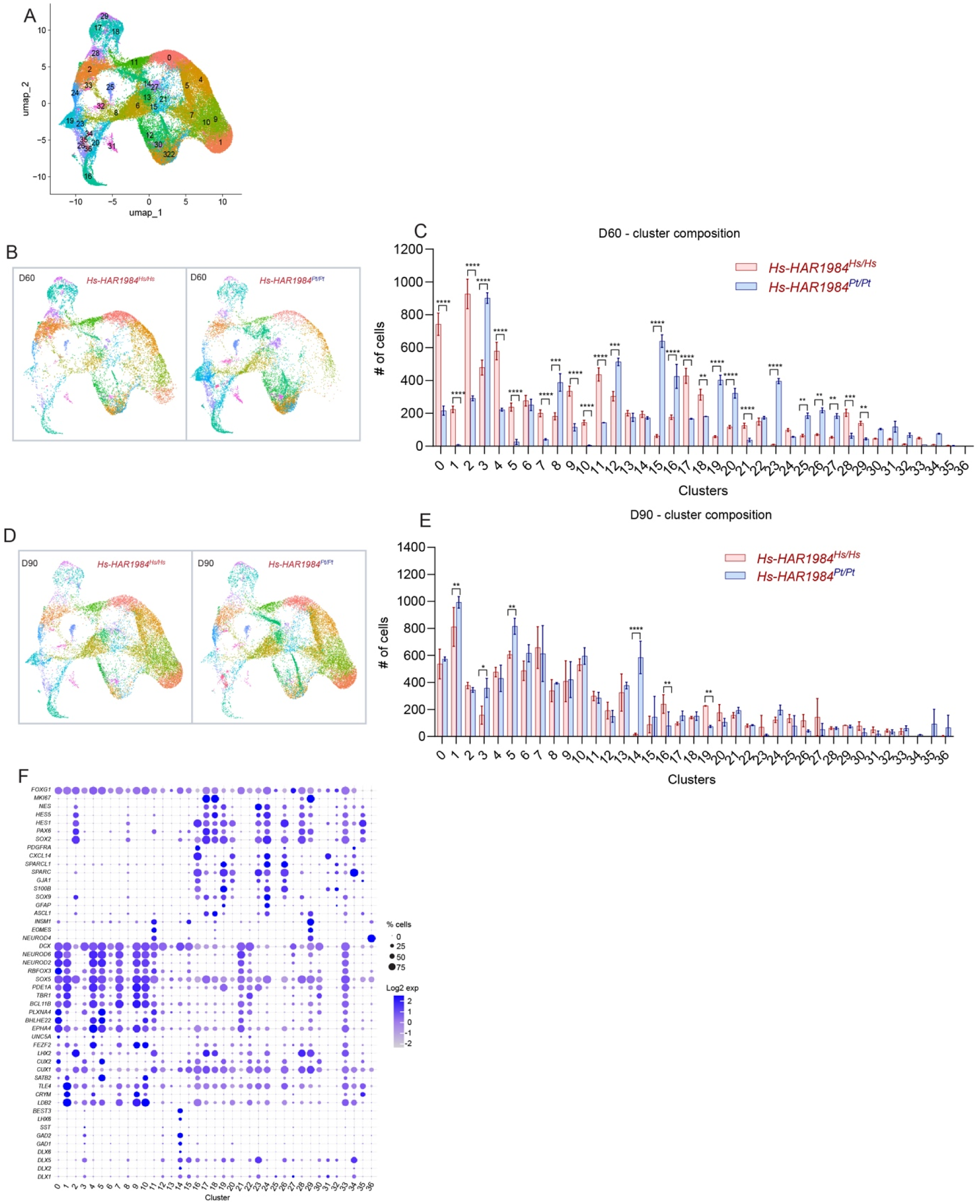
*Hs-HAR1984* impacts scRNAseq cluster composition in D60 and D90 human organoids, related to Figure 3. (A) UMAP showing integrated scRNAseq dataset divided by Seurat clusters. (B) UMAP of Seurat clusters divided by genotype at D60. (C) Number of cells in each Seurat cluster at D60. (D) UMAP of Seurat clusters divided by genotype at D90. (E) Number of cells in each Seurat cluster at D90. (F) Dot plot of top markers used to assign cell identity to Seurat clusters. Unpaired t-test (C-E). ns, not significant; \**p<*0.01; \*\**p<*0.001; \*\*\**p<*0.0001; \*\*\*\**p<*0.0001. Data are mean ± SD.

**Figure S4.**
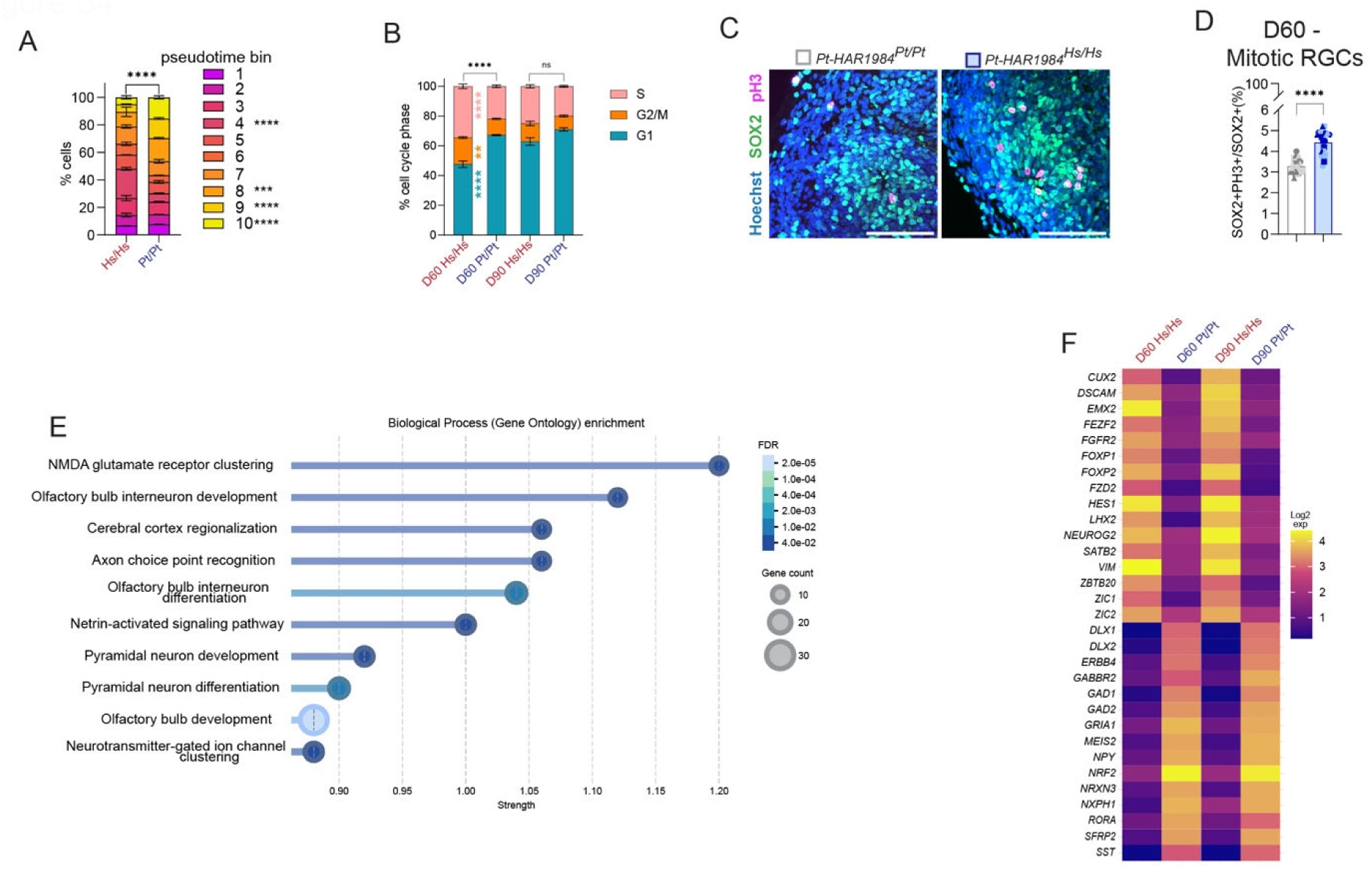
*Hs-HAR1984* promotes RGC proliferation and cortical marker expression, related to Figure 3. (A) Percentage of cells divided by pseudotime bin trajectories: from less differentiated (magenta) to more differentiated (yellow). (B) Stack histogram of scRNAseq dataset cell cycle composition. (C) Immunostaining for SOX2 (RGCs, green) and PH3 (mitotic cells, magenta) in D60 chimpanzee organoids. (D) Quantification of SOX2+PH3+ percentage over total number of RGCs (SOX2) shown in C. 3 differentiations, 3-4 organoids per differentiations. (E) GO analysis of DEGs in common between D60 and D90 in KI organoids compared to control. (F) Examples of DEGs upregulated or downregulated in KI organoids. Scale bars: 100μm. Each dot represents a single organoid. Different dot shapes represent different batches of differentiation. Unpaired t-test (D). Two-way ANOVA with Šídák correction (A, B). ns, not significant; \**p<*0.01; \*\**p<*0.001; \*\*\**p<*0.0001; \*\*\*\**p<*0.0001. Data are mean ± SD.

**Figure S5.**
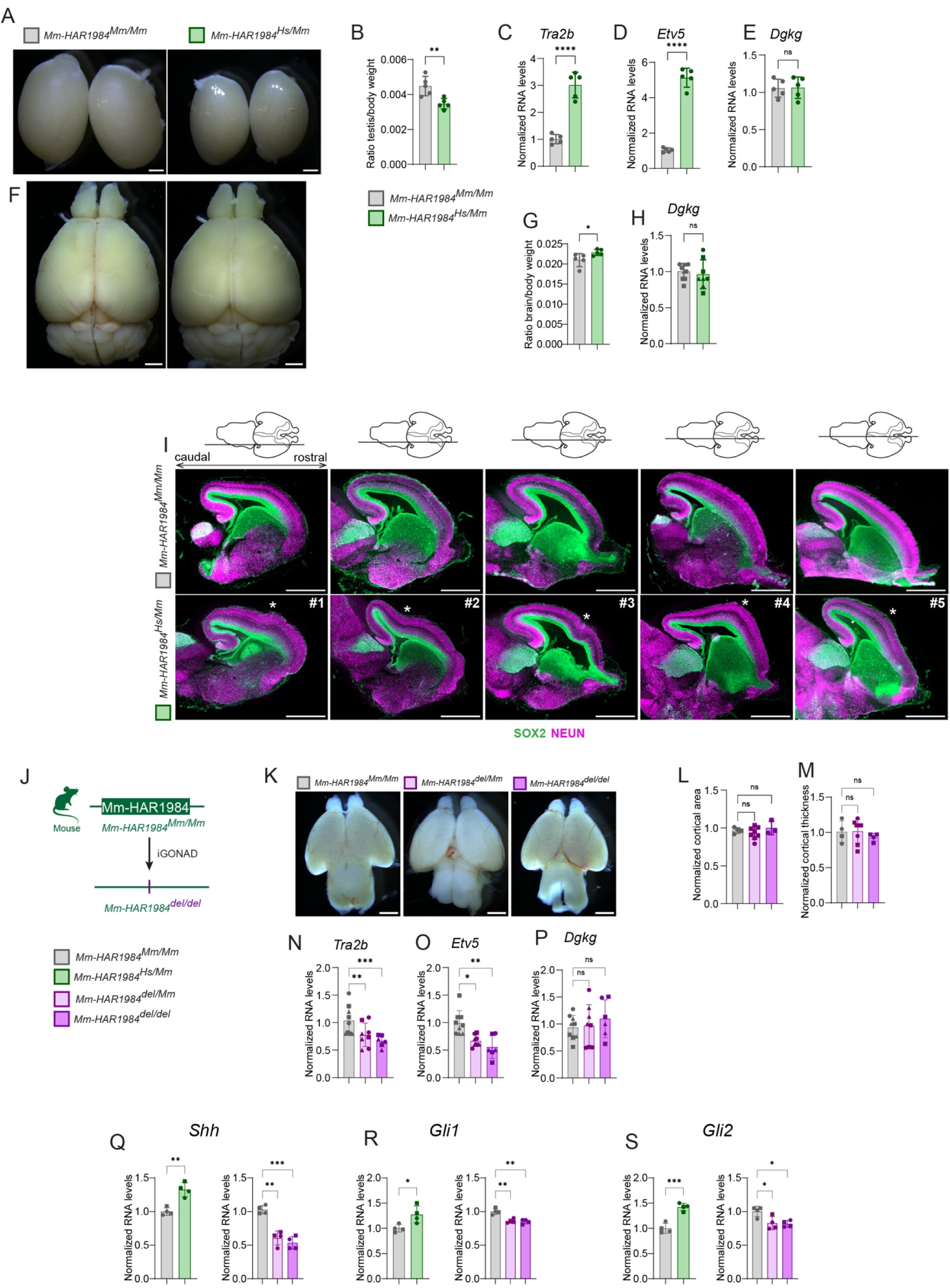
*Hs-HAR1984* promotes male fertility, brain size and cortical folding and is required for gene expression, related to Figure 4. (A) Whole mount images of testis from 12 week old control and *Mm-HAR1984^Hs/Mm^* mice. (B) Quantifications of testis weight normalized on body weight showed in A. (C-E) RT-qPCRs 12 week old mouse testis of *Tra2b* (C) and *Etv5* (D), and *Dgkg* (E). (F) Whole mount images of brain from 12 week old control and *Mm-HAR1984^Hs/Mm^* mice. (G) Quantifications of brain weight normalized on body weight showed in F. (H) RT-qPCRs of *Dgkg* expression from E16.5 cortices of control and *Mm-HAR1984^Hs/Mm^* mice. (I) Sagittal sections of E16.5 control and *HAR1984^Hs/Mm^* brains from 5 independent brains. Depiction above show section position medio-laterally. Immunostaining for SOX2 (RGCs, green) and NeuN (neurons, magenta). Asterisks indicate folding regions. (J) Scheme of iGONAD mutagenesis to generated *HAR1984* deletion mouse. (K) Whole mount images of E16.5 mouse control, *Mm-HAR1984^del/Mm^* and *Mm-HAR1984^del/del^* brains. (L-M) Quantifications of E16.5 brain area and cortical thickness in control, *Mm-HAR1984^del/Mm^* and *Mm-HAR1984^del/del^*. (N-P) RT-qPCRs E16.5 mouse brain of *Tra2b* (N) and *Etv5* (O), and *Dgkg* (P) in control, *Mm-HAR1984^del/Mm^* and *Mm-HAR1984^del/del^*. (Q-S) RT-qPCR validation of biologically meaningful DEGs identified in Bulk RNA-seq dataset in control, *HAR1984* humanized and deletion E16.5 mouse brains. Scale bars: 1mm (A, F, I, K). Each dot represents a single embryo or mouse. Unpaired t-test (B-E, G-H, Q-S). Two-way ANOVA with Šídák correction (L-M, N-P, Q-S). ns, not significant; \**p<*0.01; \*\**p<*0.001; \*\*\**p<*0.0001; \*\*\*\**p<*0.0001. Data are mean ± SD.

**Figure S6.**
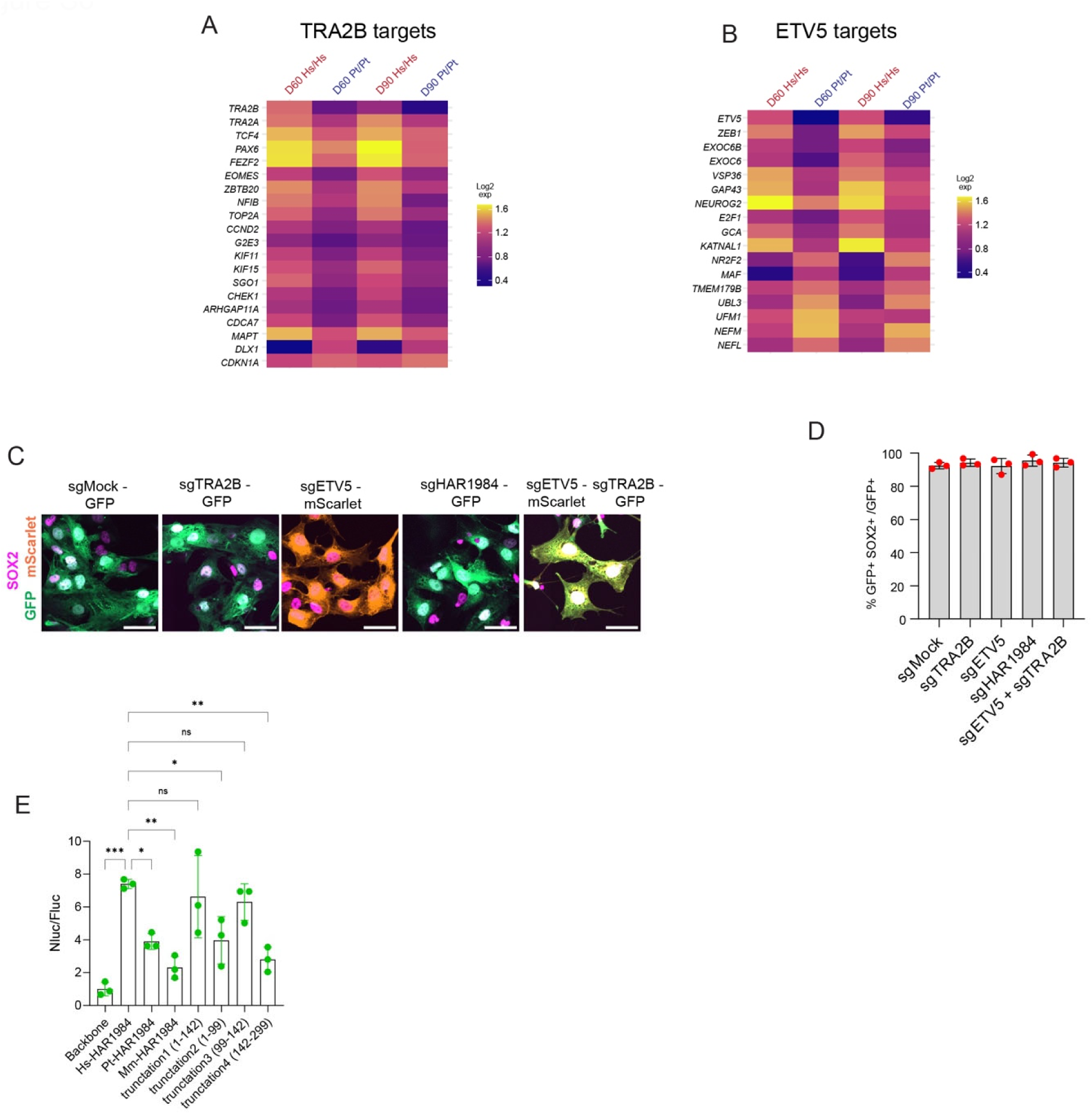
ETV5 and TRA2B targets are differentially expressed in human cortical organoids, related to Figure 5. (A-B) Heatmap of known TRA2B (A) and ETV5 (B) targets that are differentially regulated in scRNA-seq dataset in human cortical organoids at D60 and D90. (C) Representative images from dissociated organoids after 5days of sgRNAs (green or orange) viral transduction and immunostained for SOX2 (magenta). (D) Quantifications of GFP+SOX2+ cells normalized over GFP+ cells shown in C. (E) Nanoluciferase (Nluc) levels normalized to firefly (Fluc) control in D12 NPCs. GFP was co-delivered with luciferase reporter. Each dot represents an independent differentiation batch (n=3 technical replicates for each differentiation). Scale bars: 10 μm. Each dot represents a single differentiation. Two-way ANOVA with Šídák correction. ns, not significant; \**p<*0.01; \*\**p<*0.001; \*\*\**p<*0.0001; \*\*\*\**p<*0.0001. Data are mean ± SD.

**Figure S7.**
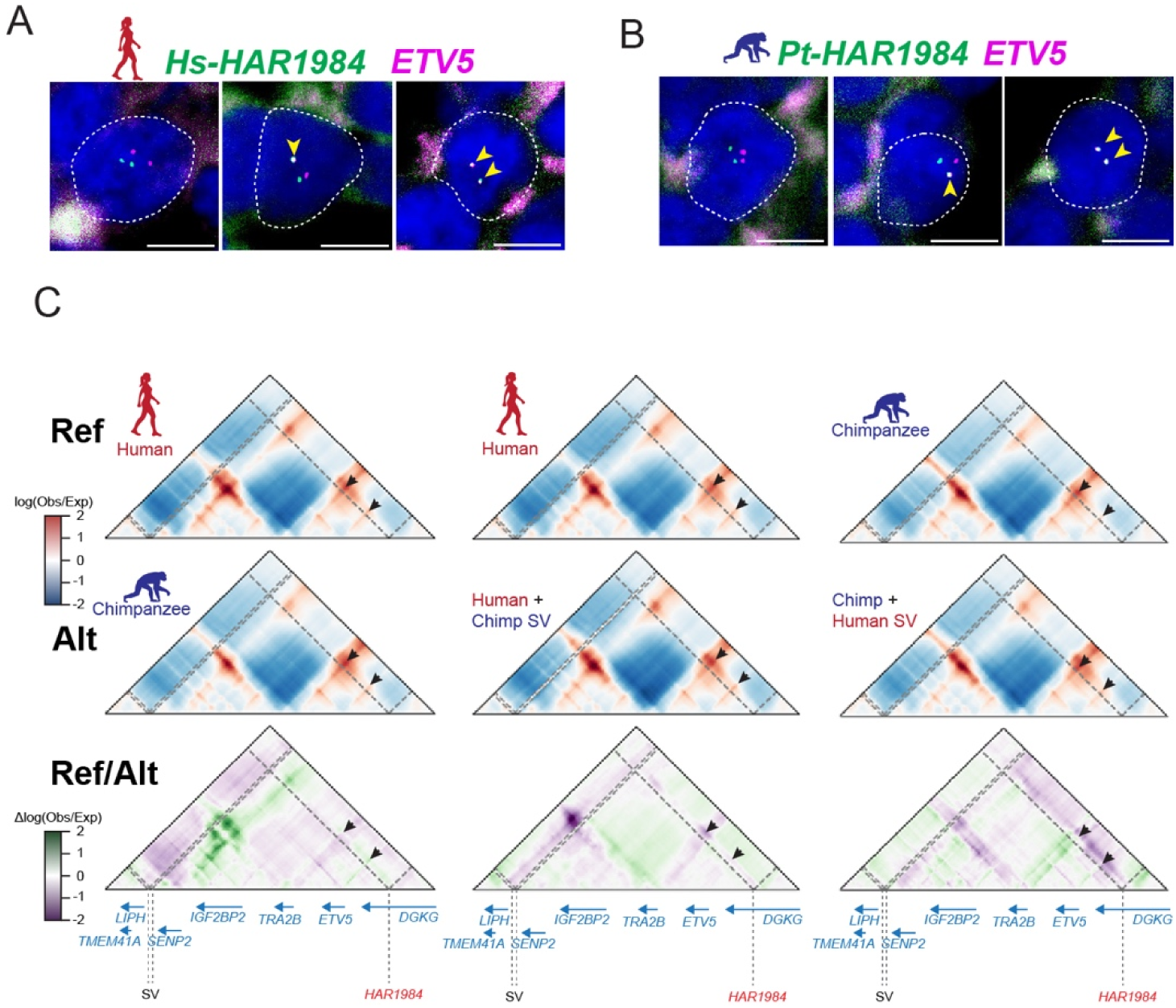
Human SVs adjacent to *Hs-HAR1984* strengthen *HAR1984*-*ETV5* and *TRA2B* promoters’ interactions, related to Figure 6. (A) Examples in human NPCs of no overlap between *HAR1984* (green) and *ETV5* (magenta) loci (left), only one chromosome overlaps (center), both chromosomes overlap (right). Yellow arrowpoint point to overlapping signal. (B) Examples in chimpanzee NPCs of no overlap between *HAR1984* (green) and *ETV5* (magenta) loci (left), only one chromosome overlaps (center), both chromosomes overlap (right). Yellow arrowpoint point to overlapping signal. (C) Hi-C maps of human or chimpanzee prediction with human SV365814. Ref (first row): human; human; and chimpanzee genome. Alt (second row): chimpanzee; human genome with chimpanzee SVs; chimpanzee genome with human SVs. Differences between Ref-Alt (third row): Human vs chimpanzee genomes; human vs human with chimpanzee SVs; and chimpanzee vs chimpanzee with human SVs.

**Table S1. *In silico* screen to identify HARs involved in neurodevelopment, related to Figure 1**.

**Table S2. DEGs from scRNA-seq D60 and D90 *HAR1984* human organoids, related to Figure 3**.

**Table S3. DEGs from Bulk-RNA-seq *HAR1984* humanized and deletion mice, related to Figure 4**.

**Table S4. HAR1984 transcription factor prediction and human-specific structural variations, related to Figure 5 and 6**.

**Table S5. Oligos used in this study, see methods.**

## Acknowledgements

We thank the Silver Laboratory past and present members for helpful discussions and Victor Borrell for reading the manuscript. We are grateful for the non-human primate biological materials provided by the Emory National Primate Research Center, formerly known as the Yerkes National Primate Research Center. The chimpanzee iPS cell line was a gift from Y. Gilad (funded by the Emory National Primate Research Center grant no. ORIP/OD P51OD011132). We thank Josh Huang for *Tbr2*-2A-Flp knock-in and Flp-dependent Ai65-Frt mice and Dmitry Velmeshev for sharing his sc-seq equipment. We thank the Duke University Mouse Transgenic Facility for mouse generation and maintenance, Duke University Light Microscopy Core Facility for the use of shared equipment, Duke University Sequencing and Genomic Technologies core for RNAseq and scRNA-seq library sequencing, and Duke Regeneromics core for bioinformatics. The following funding supported this study: NIH R01NS120667, R37NS110388 and R01MH132089 (to D.L.S.), Duke University Tricem Award (to F.M.).

## Contributions

F.M. and D.L.S. conceived of and designed the study. D.L.S. supervised the experiments. F.M. performed in silico screening, luciferase and LacZ experiments. F.M. and J.L. performed CRISPR editing on human and chimpanzee ES cells/iPS cells. F.M. performed cortical organoid experiments with the help of J.L. and V.A.K. for data analysis. F.M. performed scRNA-seq and analysis. S.S. and N.S.H. performed iGONAD mutagenesis. F.M., J.L. and K.L. performed the experiments and analyses of the mouse model. F.M. performed RNA-seq and analysis and DNA FISH experiments. K.G and K.S.P. performed chromatin interaction prediction and modeling. F.M. and D.L.S. wrote the paper. All authors reviewed, edited and approved the paper.

## Notes

### Competing Interest Statement

The authors have declared no competing interest.

## References

Aldea, D., Atsuta, Y., Kokalari, B., Schaffner, S. F., Prasasya, R. D., Aharoni, A., Dingwall, H. L., Warder, B., & Kamberov, Y. G. (2021). Repeated mutation of a developmental enhancer contributed to human thermoregulatory evolution. Proc Natl Acad Sci U S A, 118(16). 10.1073/pnas.2021722118

Amberg, N., Laukoter, S., & Hippenmeyer, S. (2019). Epigenetic cues modulating the generation of cell-type diversity in the cerebral cortex. J Neurochem, 149(1), 12–26. 10.1111/jnc.14601

Anderson, E. S., Lin, C. H., Xiao, X., Stoilov, P., Burge, C. B., & Black, D. L. (2012). The cardiotonic steroid digitoxin regulates alternative splicing through depletion of the splicing factors SRSF3 and TRA2B. Rna, 18(5), 1041–1049. 10.1261/rna.032912.112

Bankhead, P., Loughrey, M. B., Fernández, J. A., Dombrowski, Y., McArt, D. G., Dunne, P. D., McQuaid, S., Gray, R. T., Murray, L. J., Coleman, H. G., James, J. A., Salto-Tellez, M., & Hamilton, P. W. (2017). QuPath: Open source software for digital pathology image analysis. Sci Rep, 7(1), 16878. 10.1038/s41598-017-17204-5

Bayani, J., & Squire, J. A. (2004). Fluorescence in situ Hybridization (FISH). Curr Protoc Cell Biol, Chapter 22, Unit 22.24. 10.1002/0471143030.cb2204s23

Beagrie, R. A., Scialdone, A., Schueler, M., Kraemer, D. C., Chotalia, M., Xie, S. Q., Barbieri, M., de Santiago, I., Lavitas, L. M., Branco, M. R., Fraser, J., Dostie, J., Game, L., Dillon, N., Edwards, P. A., Nicodemi, M., & Pombo, A. (2017). Complex multi-enhancer contacts captured by genome architecture mapping. Nature, 543(7646), 519–524. 10.1038/nature21411

Best, A., James, K., Dalgliesh, C., Hong, E., Kheirolahi-Kouhestani, M., Curk, T., Xu, Y., Danilenko, M., Hussain, R., Keavney, B., Wipat, A., Klinck, R., Cowell, I. G., Cheong Lee, K., Austin, C. A., Venables, J. P., Chabot, B., Santibanez Koref, M., Tyson-Capper, A., & Elliott, D. J. (2014). Human Tra2 proteins jointly control a CHEK1 splicing switch among alternative and constitutive target exons. Nat Commun, 5, 4760. 10.1038/ncomms5760

Bird, C. P., Stranger, B. E., Liu, M., Thomas, D. J., Ingle, C. E., Beazley, C., Miller, W., Hurles, M. E., & Dermitzakis, E. T. (2007). Fast-evolving noncoding sequences in the human genome. Genome Biol, 8(6), R118. 10.1186/gb-2007-8-6-r118

Borrell, V., & Götz, M. (2014). Role of radial glial cells in cerebral cortex folding. Curr Opin Neurobiol, 27, 39–46. 10.1016/j.conb.2014.02.007

Boyd, J. L., Skove, S. L., Rouanet, J. P., Pilaz, L. J., Bepler, T., Gordân, R., Wray, G. A., & Silver, D. L. (2015). Human-chimpanzee differences in a FZD8 enhancer alter cell-cycle dynamics in the developing neocortex. Curr Biol, 25(6), 772–779. 10.1016/j.cub.2015.01.041

Bush, E. C., & Lahn, B. T. (2008). A genome-wide screen for noncoding elements important in primate evolution. BMC Evol Biol, 8, 17. 10.1186/1471-2148-8-17

Cai, E., Barba, M. G., & Ge, X. (2023). Hedgehog Signaling in Cortical Development. Cells, 13(1). 10.3390/cells13010021

Capra, J. A., Erwin, G. D., McKinsey, G., Rubenstein, J. L., & Pollard, K. S. (2013). Many human accelerated regions are developmental enhancers. Philos Trans R Soc Lond B Biol Sci, 368(1632), 20130025. 10.1098/rstb.2013.0025

Cárdenas, A., Villalba, A., de Juan Romero, C., Picó, E., Kyrousi, C., Tzika, A. C., Tessier-Lavigne, M., Ma, L., Drukker, M., Cappello, S., & Borrell, V. (2018). Evolution of Cortical Neurogenesis in Amniotes Controlled by Robo Signaling Levels. Cell, 174(3), 590–606.e521. 10.1016/j.cell.2018.06.007

Cecchi, C., & Boncinelli, E. (2000). Emx homeogenes and mouse brain development. Trends Neurosci, 23(8), 347–352. 10.1016/s0166-2236(00)01608-8

Charrier, C., Joshi, K., Coutinho-Budd, J., Kim, J. E., Lambert, N., de Marchena, J., Jin, W. L., Vanderhaeghen, P., Ghosh, A., Sassa, T., & Polleux, F. (2012). Inhibition of SRGAP2 function by its human-specific paralogs induces neoteny during spine maturation. Cell, 149(4), 923–935. 10.1016/j.cell.2012.03.034

Chen, H., Li, C., Zhou, Z., & Liang, H. (2018). Fast-Evolving Human-Specific Neural Enhancers Are Associated with Aging-Related Diseases. Cell Syst, 6(5), 604–611.e604. 10.1016/j.cels.2018.04.002

Choudhary, M. N. K., Quaid, K., Xing, X., Schmidt, H., & Wang, T. (2023). Widespread contribution of transposable elements to the rewiring of mammalian 3D genomes. Nat Commun, 14(1), 634. 10.1038/s41467-023-36364-9

Co, M., Anderson, A. G., & Konopka, G. (2020). FOXP transcription factors in vertebrate brain development, function, and disorders. Wiley Interdiscip Rev Dev Biol, 9(5), e375. 10.1002/wdev.375

Concordet, J. P., & Haeussler, M. (2018). CRISPOR: intuitive guide selection for CRISPR/Cas9 genome editing experiments and screens. Nucleic Acids Res, 46(W1), W242–w245. 10.1093/nar/gky354

Cui, X., Yang, H., Cai, C., Beaman, C., Yang, X., Liu, H., Ren, X., Amador, Z., Jones, I. R., Keough, K. C., Zhang, M., Fair, T., Abnousi, A., Mishra, S., Ye, Z., Hu, M., Pollen, A. A., Pollard, K. S., & Shen, Y. (2025). Comparative characterization of human accelerated regions in neurons. Nature, 640(8060), 991–999. 10.1038/s41586-025-08622-x

Dalgliesh, C., Aldalaqan, S., Atallah, C., Best, A., Scott, E., Ehrmann, I., Merces, G., Mannion, J., Badurova, B., Sandher, R., Illing, Y., Wirth, B., Wells, S., Codner, G., Teboul, L., Smith, G. R., Hedley, A., Herbert, M., de Rooij, D. G., … Elliott, D. J. (2025). An ultra-conserved poison exon in the Tra2b gene encoding a splicing activator is essential for male fertility and meiotic cell division. Embo j, 44(3), 877–902. 10.1038/s44318-024-00344-6

Dave, R. K., Ellis, T., Toumpas, M. C., Robson, J. P., Julian, E., Adolphe, C., Bartlett, P. F., Cooper, H. M., Reynolds, B. A., & Wainwright, B. J. (2011). Sonic hedgehog and notch signaling can cooperate to regulate neurogenic divisions of neocortical progenitors. PLoS One, 6(2), e14680. 10.1371/journal.pone.0014680

Del Toro, D., Ruff, T., Cederfjäll, E., Villalba, A., Seyit-Bremer, G., Borrell, V., & Klein, R. (2017). Regulation of Cerebral Cortex Folding by Controlling Neuronal Migration via FLRT Adhesion Molecules. Cell, 169(4), 621–635.e616. 10.1016/j.cell.2017.04.012

Del-Valle-Anton, L., Amin, S., Cimino, D., Neuhaus, F., Dvoretskova, E., Fernández, V., Babal, Y. K., Garcia-Frigola, C., Prieto-Colomina, A., Murcia-Ramón, R., Nomura, Y., Cárdenas, A., Feng, C., Moreno-Bravo, J. A., Götz, M., Mayer, C., & Borrell, V. (2024). Multiple parallel cell lineages in the developing mammalian cerebral cortex. Sci Adv, 10(13), eadn9998. 10.1126/sciadv.adn9998

Di Bella, D. J., Habibi, E., Stickels, R. R., Scalia, G., Brown, J., Yadollahpour, P., Yang, S. M., Abbate, C., Biancalani, T., Macosko, E. Z., Chen, F., Regev, A., & Arlotta, P. (2021). Molecular logic of cellular diversification in the mouse cerebral cortex. Nature, 595(7868), 554–559. 10.1038/s41586-021-03670-5

Doan, R. N., Bae, B. I., Cubelos, B., Chang, C., Hossain, A. A., Al-Saad, S., Mukaddes, N. M., Oner, O., Al-Saffar, M., Balkhy, S., Gascon, G. G., Nieto, M., & Walsh, C. A. (2016). Mutations in Human Accelerated Regions Disrupt Cognition and Social Behavior. Cell, 167(2), 341–354.e312. 10.1016/j.cell.2016.08.071

Dobin, A., Davis, C. A., Schlesinger, F., Drenkow, J., Zaleski, C., Jha, S., Batut, P., Chaisson, M., & Gingeras, T. R. (2013). STAR: ultrafast universal RNA-seq aligner. Bioinformatics, 29(1), 15–21. 10.1093/bioinformatics/bts635

Dutrow, E. V., Emera, D., Yim, K., Uebbing, S., Kocher, A. A., Krenzer, M., Nottoli, T., Burkhardt, D. B., Krishnaswamy, S., Louvi, A., & Noonan, J. P. (2022). Modeling uniquely human gene regulatory function via targeted humanization of the mouse genome. Nat Commun, 13(1), 304. 10.1038/s41467-021-27899-w

Fiddes, I. T., Lodewijk, G. A., Mooring, M., Bosworth, C. M., Ewing, A. D., Mantalas, G. L., Novak, A. M., van den Bout, A., Bishara, A., Rosenkrantz, J. L., Lorig-Roach, R., Field, A. R., Haeussler, M., Russo, L., Bhaduri, A., Nowakowski, T. J., Pollen, A. A., Dougherty, M. L., Nuttle, X., … Haussler, D. (2018). Human-Specific NOTCH2NL Genes Affect Notch Signaling and Cortical Neurogenesis. Cell, 173(6), 1356–1369.e1322. 10.1016/j.cell.2018.03.051

Fiddes, I. T., Pollen, A. A., Davis, J. M., & Sikela, J. M. (2019). Paired involvement of human-specific Olduvai domains and NOTCH2NL genes in human brain evolution. Hum Genet, 138(7), 715–721. 10.1007/s00439-019-02018-4

Fietz, S. A., Lachmann, R., Brandl, H., Kircher, M., Samusik, N., Schröder, R., Lakshmanaperumal, N., Henry, I., Vogt, J., Riehn, A., Distler, W., Nitsch, R., Enard, W., Pääbo, S., & Huttner, W. B. (2012). Transcriptomes of germinal zones of human and mouse fetal neocortex suggest a role of extracellular matrix in progenitor self-renewal. Proc Natl Acad Sci U S A, 109(29), 11836–11841. 10.1073/pnas.1209647109

Fischer, J., Fernández Ortuño, E., Marsoner, F., Artioli, A., Peters, J., Namba, T., Eugster Oegema, C., Huttner, W. B., Ladewig, J., & Heide, M. (2022). Human-specific ARHGAP11B ensures human-like basal progenitor levels in hominid cerebral organoids. EMBO Rep, 23(11), e54728. 10.15252/embr.202254728

Florio, M., Albert, M., Taverna, E., Namba, T., Brandl, H., Lewitus, E., Haffner, C., Sykes, A., Wong, F. K., Peters, J., Guhr, E., Klemroth, S., Prüfer, K., Kelso, J., Naumann, R., Nüsslein, I., Dahl, A., Lachmann, R., Pääbo, S., & Huttner, W. B. (2015). Human-specific gene ARHGAP11B promotes basal progenitor amplification and neocortex expansion. Science, 347(6229), 1465–1470. 10.1126/science.aaa1975

Florio, M., Heide, M., Pinson, A., Brandl, H., Albert, M., Winkler, S., Wimberger, P., Huttner, W. B., & Hiller, M. (2018). Evolution and cell-type specificity of human-specific genes preferentially expressed in progenitors of fetal neocortex. Elife, 7. 10.7554/eLife.32332

Fontanet, P. A., Ríos, A. S., Alsina, F. C., Paratcha, G., & Ledda, F. (2018). Pea3 Transcription Factors, Etv4 and Etv5, Are Required for Proper Hippocampal Dendrite Development and Plasticity. Cereb Cortex, 28(1), 236–249. 10.1093/cercor/bhw372

Franke, M., Ibrahim, D. M., Andrey, G., Schwarzer, W., Heinrich, V., Schöpflin, R., Kraft, K., Kempfer, R., Jerković, I., Chan, W. L., Spielmann, M., Timmermann, B., Wittler, L., Kurth, I., Cambiaso, P., Zuffardi, O., Houge, G., Lambie, L., Brancati, F., … Mundlos, S. (2016). Formation of new chromatin domains determines pathogenicity of genomic duplications. Nature, 538(7624), 265–269. 10.1038/nature19800

Fudenberg, G., Kelley, D. R., & Pollard, K. S. (2020). Predicting 3D genome folding from DNA sequence with Akita. Nat Methods, 17(11), 1111–1117. 10.1038/s41592-020-0958-x

Gallego Romero, I., Pavlovic, B. J., Hernando-Herraez, I., Zhou, X., Ward, M. C., Banovich, N. E., Kagan, C. L., Burnett, J. E., Huang, C. H., Mitrano, A., Chavarria, C. I., Friedrich Ben-Nun, I., Li, Y., Sabatini, K., Leonardo, T. R., Parast, M., Marques-Bonet, T., Laurent, L. C., Loring, J. F., & Gilad, Y. (2015). A panel of induced pluripotent stem cells from chimpanzees: a resource for comparative functional genomics. Elife, 4, e07103. 10.7554/eLife.07103

Geschwind, D. H., & Rakic, P. (2013). Cortical evolution: judge the brain by its cover. Neuron, 80(3), 633–647. 10.1016/j.neuron.2013.10.045

Girskis, K. M., Stergachis, Andrew B., Degennaro, Ellen M., Doan, R. N., Qian, X., Johnson, M. B., Wang, P. P., Sejourne, G. M., Nagy, M. A., Pollina, E. A., Sousa, A. M. M., Shin, T., Kenny, C. J., Scotellaro, J. L., Debo, B. M., Gonzalez, D. M., Rento, L. M., Yeh, R. C., Song, J. H. T., Beaudin, M., Fan, J., … Walsh, C. A. (2021). Rewiring of human neurodevelopmental gene regulatory programs by human accelerated regions. Neuron. 10.1016/j.neuron.2021.08.005

Gittelman, R. M., Hun, E., Ay, F., Madeoy, J., Pennacchio, L., Noble, W. S., Hawkins, R. D., & Akey, J. M. (2015). Comprehensive identification and analysis of human accelerated regulatory DNA. Genome Res, 25(9), 1245–1255. 10.1101/gr.192591.115

Gjoni, K., & Pollard, K. S. (2024). SuPreMo: a computational tool for streamlining in silico perturbation using sequence-based predictive models. Bioinformatics, 40(6). 10.1093/bioinformatics/btae340

Gokhman, D., Agoglia, R. M., Kinnebrew, M., Gordon, W., Sun, D., Bajpai, V. K., Naqvi, S., Chen, C., Chan, A., Chen, C., Petrov, D. A., Ahituv, N., Zhang, H., Mishina, Y., Wysocka, J., Rohatgi, R., & Fraser, H. B. (2021). Human-chimpanzee fused cells reveal cis-regulatory divergence underlying skeletal evolution. Nat Genet, 53(4), 467–476. 10.1038/s41588-021-00804-3

Gorkin, D. U., Barozzi, I., Zhao, Y., Zhang, Y., Huang, H., Lee, A. Y., Li, B., Chiou, J., Wildberg, A., Ding, B., Zhang, B., Wang, M., Strattan, J. S., Davidson, J. M., Qiu, Y., Afzal, V., Akiyama, J. A., Plajzer-Frick, I., Novak, C. S., … Ren, B. (2020). An atlas of dynamic chromatin landscapes in mouse fetal development. Nature, 583(7818), 744–751. 10.1038/s41586-020-2093-3

Grellscheid, S., Dalgliesh, C., Storbeck, M., Best, A., Liu, Y., Jakubik, M., Mende, Y., Ehrmann, I., Curk, T., Rossbach, K., Bourgeois, C. F., Stévenin, J., Grellscheid, D., Jackson, M. S., Wirth, B., & Elliott, D. J. (2011). Identification of evolutionarily conserved exons as regulated targets for the splicing activator tra2β in development. PLoS Genet, 7(12), e1002390. 10.1371/journal.pgen.1002390

Gurumurthy, C. B., Sato, M., Nakamura, A., Inui, M., Kawano, N., Islam, M. A., Ogiwara, S., Takabayashi, S., Matsuyama, M., Nakagawa, S., Miura, H., & Ohtsuka, M. (2019). Creation of CRISPR-based germline-genome-engineered mice without ex vivo handling of zygotes by i-GONAD. Nat Protoc, 14(8), 2452–2482. 10.1038/s41596-019-0187-x

Harabula, I., & Pombo, A. (2021). The dynamics of chromatin architecture in brain development and function. Curr Opin Genet Dev, 67, 84–93. 10.1016/j.gde.2020.12.008

Heide, M., Haffner, C., Murayama, A., Kurotaki, Y., Shinohara, H., Okano, H., Sasaki, E., & Huttner, W. B. (2020). Human-specific ARHGAP11B increases size and folding of primate neocortex in the fetal marmoset. Science, 369(6503), 546–550. 10.1126/science.abb2401

Heide, M., & Huttner, W. B. (2021). Human-Specific Genes, Cortical Progenitor Cells, and Microcephaly. Cells, 10(5). 10.3390/cells10051209

Hou, S., Ho, W. L., Wang, L., Kuo, B., Park, J. Y., & Han, Y. G. (2021). Biphasic Roles of Hedgehog Signaling in the Production and Self-Renewal of Outer Radial Glia in the Ferret Cerebral Cortex. Cereb Cortex, 31(10), 4730–4741. 10.1093/cercor/bhab119

Huilgol, D., Russ, J. B., Srivas, S., & Huang, Z. J. (2023). The progenitor basis of cortical projection neuron diversity. Curr Opin Neurobiol, 81, 102726. 10.1016/j.conb.2023.102726

Javier-Torrent, M., Zimmer-Bensch, G., & Nguyen, L. (2021). Mechanical Forces Orchestrate Brain Development. Trends Neurosci, 44(2), 110–121. 10.1016/j.tins.2020.10.012

Johnson, M. B., Wang, P. P., Atabay, K. D., Murphy, E. A., Doan, R. N., Hecht, J. L., & Walsh, C. A. (2015). Single-cell analysis reveals transcriptional heterogeneity of neural progenitors in human cortex. Nat Neurosci, 18(5), 637–646. 10.1038/nn.3980

Jorstad, N. L., Song, J. H. T., Exposito-Alonso, D., Suresh, H., Castro-Pacheco, N., Krienen, F. M., Yanny, A. M., Close, J., Gelfand, E., Long, B., Seeman, S. C., Travaglini, K. J., Basu, S., Beaudin, M., Bertagnolli, D., Crow, M., Ding, S. L., Eggermont, J., Glandon, A., … Bakken, T. E. (2023). Comparative transcriptomics reveals human-specific cortical features. Science, 382(6667), eade9516. 10.1126/science.ade9516

Jung, I., Schmitt, A., Diao, Y., Lee, A. J., Liu, T., Yang, D., Tan, C., Eom, J., Chan, M., Chee, S., Chiang, Z., Kim, C., Masliah, E., Barr, C. L., Li, B., Kuan, S., Kim, D., & Ren, B. (2019). A compendium of promoter-centered long-range chromatin interactions in the human genome. Nat Genet, 51(10), 1442–1449. 10.1038/s41588-019-0494-8

Kalkan, T., Bornelöv, S., Mulas, C., Diamanti, E., Lohoff, T., Ralser, M., Middelkamp, S., Lombard, P., Nichols, J., & Smith, A. (2019). Complementary Activity of ETV5, RBPJ, and TCF3 Drives Formative Transition from Naive Pluripotency. Cell Stem Cell, 24(5), 785–801.e787. 10.1016/j.stem.2019.03.017

Keough, K. C., Whalen, S., Inoue, F., Przytycki, P. F., Fair, T., Deng, C., Steyert, M., Ryu, H., Lindblad-Toh, K., Karlsson, E., Nowakowski, T., Ahituv, N., Pollen, A., & Pollard, K. S. (2023). Three-dimensional genome rewiring in loci with human accelerated regions. Science, 380(6643), eabm1696. 10.1126/science.abm1696

Komada, M., Saitsu, H., Kinboshi, M., Miura, T., Shiota, K., & Ishibashi, M. (2008). Hedgehog signaling is involved in development of the neocortex. Development, 135(16), 2717–2727. 10.1242/dev.015891

Kronenberg, Z. N., Fiddes, I. T., Gordon, D., Murali, S., Cantsilieris, S., Meyerson, O. S., Underwood, J. G., Nelson, B. J., Chaisson, M. J. P., Dougherty, M. L., Munson, K. M., Hastie, A. R., Diekhans, M., Hormozdiari, F., Lorusso, N., Hoekzema, K., Qiu, R., Clark, K., Raja, A., … Eichler, E. E. (2018). High-resolution comparative analysis of great ape genomes. Science, 360(6393). 10.1126/science.aar6343

Kundaje, A., Meuleman, W., Ernst, J., Bilenky, M., Yen, A., Heravi-Moussavi, A., Kheradpour, P., Zhang, Z., Wang, J., Ziller, M. J., Amin, V., Whitaker, J. W., Schultz, M. D., Ward, L. D., Sarkar, A., Quon, G., Sandstrom, R. S., Eaton, M. L., Wu, Y. C., … Kellis, M. (2015). Integrative analysis of 111 reference human epigenomes. Nature, 518(7539), 317–330. 10.1038/nature14248

Lindblad-Toh, K., Garber, M., Zuk, O., Lin, M. F., Parker, B. J., Washietl, S., Kheradpour, P., Ernst, J., Jordan, G., Mauceli, E., Ward, L. D., Lowe, C. B., Holloway, A. K., Clamp, M., Gnerre, S., Alföldi, J., Beal, K., Chang, J., Clawson, H., … Kellis, M. (2011). A high-resolution map of human evolutionary constraint using 29 mammals. Nature, 478(7370), 476–482. 10.1038/nature10530

Liu, J., Mosti, F., Zhao, H. T., Lollis, D., Sotelo-Fonseca, J. E., Escobar-Tomlienovich, C. F., Musso, C. M., Mao, Y., Massri, A. J., Doll, H. M., Moss, N. D., Sousa, A. M. M., Wray, G. A., Schmidt, E. R. E., & Silver, D. L. (2025). A human-specific enhancer fine-tunes radial glia potency and corticogenesis. Nature. 10.1038/s41586-025-09002-1

Liu, Y., & Zhang, Y. (2019). ETV5 is Essential for Neuronal Differentiation of Human Neural Progenitor Cells by Repressing NEUROG2 Expression. Stem Cell Rev Rep, 15(5), 703–716. 10.1007/s12015-019-09904-4

Llinares-Benadero, C., & Borrell, V. (2019). Deconstructing cortical folding: genetic, cellular and mechanical determinants. Nat Rev Neurosci, 20(3), 161–176. 10.1038/s41583-018-0112-2

Lodato, S., & Arlotta, P. (2015). Generating neuronal diversity in the mammalian cerebral cortex. Annu Rev Cell Dev Biol, 31, 699–720. 10.1146/annurev-cellbio-100814-125353

Lu, C., Garipler, G., Dai, C., Roush, T., Salome-Correa, J., Martin, A., Liscovitch-Brauer, N., Mazzoni, E. O., & Sanjana, N. E. (2023). Essential transcription factors for induced neuron differentiation. Nat Commun, 14(1), 8362. 10.1038/s41467-023-43602-7

Lun, A. T., McCarthy, D. J., & Marioni, J. C. (2016). A step-by-step workflow for low-level analysis of single-cell RNA-seq data with Bioconductor. F1000Res, 5, 2122. 10.12688/f1000research.9501.2

Luo, X., Liu, Y., Dang, D., Hu, T., Hou, Y., Meng, X., Zhang, F., Li, T., Wang, C., Li, M., Wu, H., Shen, Q., Hu, Y., Zeng, X., He, X., Yan, L., Zhang, S., Li, C., & Su, B. (2021). 3D Genome of macaque fetal brain reveals evolutionary innovations during primate corticogenesis. Cell, 184(3), 723–740.e721. 10.1016/j.cell.2021.01.001

Martin, V., Zhao, J., Afek, A., Mielko, Z., & Gordân, R. (2019). QBiC-Pred: quantitative predictions of transcription factor binding changes due to sequence variants. Nucleic Acids Res, 47(W1), W127–w135. 10.1093/nar/gkz363

Martínez-Cerdeño, V., Noctor, S. C., & Kriegstein, A. R. (2006). The role of intermediate progenitor cells in the evolutionary expansion of the cerebral cortex. Cereb Cortex, 16 Suppl 1, i152–161. 10.1093/cercor/bhk017

Matsumoto, N., Tanaka, S., Horiike, T., Shinmyo, Y., & Kawasaki, H. (2020). A discrete subtype of neural progenitor crucial for cortical folding in the gyrencephalic mammalian brain. Elife, 9. 10.7554/eLife.54873

Miller, E. E., Kobayashi, G. S., Musso, C. M., Allen, M., Ishiy, F. A. A., de Caires, L. C., Jr., Goulart, E., Griesi-Oliveira, K., Zechi-Ceide, R. M., Richieri-Costa, A., Bertola, D. R., Passos-Bueno, M. R., & Silver, D. L. (2017). EIF4A3 deficient human iPSCs and mouse models demonstrate neural crest defects that underlie Richieri-Costa-Pereira syndrome. Hum Mol Genet, 26(12), 2177–2191. 10.1093/hmg/ddx078

Miller, J. A., Ding, S. L., Sunkin, S. M., Smith, K. A., Ng, L., Szafer, A., Ebbert, A., Riley, Z. L., Royall, J. J., Aiona, K., Arnold, J. M., Bennet, C., Bertagnolli, D., Brouner, K., Butler, S., Caldejon, S., Carey, A., Cuhaciyan, C., Dalley, R. A., … Lein, E. S. (2014). Transcriptional landscape of the prenatal human brain. Nature, 508(7495), 199–206. 10.1038/nature13185

Miura, Y., Li, M. Y., Birey, F., Ikeda, K., Revah, O., Thete, M. V., Park, J. Y., Puno, A., Lee, S. H., Porteus, M. H., & Pașca, S. P. (2020). Generation of human striatal organoids and cortico-striatal assembloids from human pluripotent stem cells. Nat Biotechnol, 38(12), 1421–1430. 10.1038/s41587-020-00763-w

Molnar, Z., Clowry, G. J., Sestan, N., Alzu’bi, A., Bakken, T., Hevner, R. F., Huppi, P. S., Kostovic, I., Rakic, P., Anton, E. S., Edwards, D., Garcez, P., Hoerder-Suabedissen, A., & Kriegstein, A. (2019). New insights into the development of the human cerebral cortex. J Anat, 235(3), 432–451. 10.1111/joa.13055

Morrow, C. M., Hostetler, C. E., Griswold, M. D., Hofmann, M. C., Murphy, K. M., Cooke, P. S., & Hess, R. A. (2007). ETV5 is required for continuous spermatogenesis in adult mice and may mediate blood testes barrier function and testicular immune privilege. Ann N Y Acad Sci, 1120, 144–151. 10.1196/annals.1411.005

Nestorowa, S., Hamey, F. K., Pijuan Sala, B., Diamanti, E., Shepherd, M., Laurenti, E., Wilson, N. K., Kent, D. G., & Göttgens, B. (2016). A single-cell resolution map of mouse hematopoietic stem and progenitor cell differentiation. Blood, 128(8), e20–31. 10.1182/blood-2016-05-716480

Nonaka-Kinoshita, M., Reillo, I., Artegiani, B., Martínez-Martínez, M., Nelson, M., Borrell, V., & Calegari, F. (2013). Regulation of cerebral cortex size and folding by expansion of basal progenitors. Embo j, 32(13), 1817–1828. 10.1038/emboj.2013.96

Norman, A. R., Ryu, A. H., Jamieson, K., Thomas, S., Shen, Y., Ahituv, N., Pollard, K. S., & Reiter, J. F. (2021). A Human Accelerated Region is a Leydig cell GLI2 Enhancer that Affects Male-Typical Behavior. bioRxiv.

Nowakowski, T. J., Bhaduri, A., Pollen, A. A., Alvarado, B., Mostajo-Radji, M. A., Di Lullo, E., Haeussler, M., Sandoval-Espinosa, C., Liu, S. J., Velmeshev, D., Ounadjela, J. R., Shuga, J., Wang, X., Lim, D. A., West, J. A., Leyrat, A. A., Kent, W. J., & Kriegstein, A. R. (2017). Spatiotemporal gene expression trajectories reveal developmental hierarchies of the human cortex. Science, 358(6368), 1318–1323. 10.1126/science.aap8809

Pal, A., Noble, M. A., Morales, M., Pal, R., Baumgartner, M., Yang, J. W., Yim, K. M., Uebbing, S., & Noonan, J. P. (2025). Resolving the three-dimensional interactome of human accelerated regions during human and chimpanzee neurodevelopment. Cell, 188(6), 1504–1523.e1527. 10.1016/j.cell.2025.01.007

Pebworth, M. P., Ross, J., Andrews, M., Bhaduri, A., & Kriegstein, A. R. (2021). Human intermediate progenitor diversity during cortical development. Proc Natl Acad Sci U S A, 118(26). 10.1073/pnas.2019415118

Pollard, K. S., Salama, S. R., King, B., Kern, A. D., Dreszer, T., Katzman, S., Siepel, A., Pedersen, J. S., Bejerano, G., Baertsch, R., Rosenbloom, K. R., Kent, J., & Haussler, D. (2006). Forces shaping the fastest evolving regions in the human genome. PLoS Genet, 2(10), e168. 10.1371/journal.pgen.0020168

Pollen, A. A., Nowakowski, T. J., Chen, J., Retallack, H., Sandoval-Espinosa, C., Nicholas, C. R., Shuga, J., Liu, S. J., Oldham, M. C., Diaz, A., Lim, D. A., Leyrat, A. A., West, J. A., & Kriegstein, A. R. (2015). Molecular identity of human outer radial glia during cortical development. Cell, 163(1), 55–67. 10.1016/j.cell.2015.09.004

Prabhakar, S., Noonan, J. P., Pääbo, S., & Rubin, E. M. (2006). Accelerated evolution of conserved noncoding sequences in humans. Science, 314(5800), 786. 10.1126/science.1130738

Pravata, M. V., Forero, A., Ayo Martin, A. C., Berto, G., Heymann, T., Fast, L., Mann, M., Riesenberg, S., & Cappello, S. (2025). DCHS1 Modulates Forebrain Proportions in Modern Humans via a Glycosylation Change. bioRxiv. 10.1101/2025.05.14.654031

Preuss, T. M., & Wise, S. P. (2022). Evolution of prefrontal cortex. Neuropsychopharmacology, 47(1), 3–19. 10.1038/s41386-021-01076-5

Qiu, X., Mao, Q., Tang, Y., Wang, L., Chawla, R., Pliner, H. A., & Trapnell, C. (2017). Reversed graph embedding resolves complex single-cell trajectories. Nat Methods, 14(10), 979–982. 10.1038/nmeth.4402

Rilling, J. K. (2014). Comparative primate neurobiology and the evolution of brain language systems. Curr Opin Neurobiol, 28, 10–14. 10.1016/j.conb.2014.04.002

Roberts, J. M., Ennajdaoui, H., Edmondson, C., Wirth, B., Sanford, J. R., & Chen, B. (2014). Splicing factor TRA2B is required for neural progenitor survival. J Comp Neurol, 522(2), 372–392. 10.1002/cne.23405

Robinson, M. D., McCarthy, D. J., & Smyth, G. K. (2010). edgeR: a Bioconductor package for differential expression analysis of digital gene expression data. Bioinformatics, 26(1), 139–140. 10.1093/bioinformatics/btp616

Saito, T., & Nakatsuji, N. (2001). Efficient gene transfer into the embryonic mouse brain using in vivo electroporation. Dev Biol, 240(1), 237–246. 10.1006/dbio.2001.0439

Schindelin, J., Arganda-Carreras, I., Frise, E., Kaynig, V., Longair, M., Pietzsch, T., Preibisch, S., Rueden, C., Saalfeld, S., Schmid, B., Tinevez, J. Y., White, D. J., Hartenstein, V., Eliceiri, K., Tomancak, P., & Cardona, A. (2012). Fiji: an open-source platform for biological-image analysis. Nat Methods, 9(7), 676–682. 10.1038/nmeth.2019

Shi, Y., Kirwan, P., & Livesey, F. J. (2012). Directed differentiation of human pluripotent stem cells to cerebral cortex neurons and neural networks. Nature Protocols, 7(10), 1836–1846. 10.1038/nprot.2012.116

Shin, T., Song, J. H. T., Kosicki, M., Kenny, C., Beck, S. G., Kelley, L., Antony, I., Qian, X., Bonacina, J., Papandile, F., Gonzalez, D., Scotellaro, J., Bushinsky, E. M., Andersen, R. E., Maury, E., Pennacchio, L. A., Doan, R. N., & Walsh, C. A. (2024). Rare variation in non-coding regions with evolutionary signatures contributes to autism spectrum disorder risk. Cell Genom, 4(8), 100609. 10.1016/j.xgen.2024.100609

Silver, D. L., Rakic, P., Grove, E.A., Haydar, T.F., Hensch, T.K., Huttner, W.B., Molnar, z., Rubenstein, J.L., Sestan, N., Stryker, M.P., Sur, M., Tosches, M.A., Walsh, C. (2019). Evolution and ontogenetic development of cortical structures. In: The Neocortex. In s. e. S. u. F. R. J. R. Lupp (Ed.), ((eds Singer W, Sejnowski TJ, Rakic P) ed., Vol. Vol. 27). MIT Press, in press.

Singh, A., Del-Valle-Anton, L., de Juan Romero, C., Zhang, Z., Ortuño, E. F., Mahesh, A., Espinós, A., Soler, R., Cárdenas, A., Fernández, V., Lusby, R., Tiwari, V. K., & Borrell, V. (2024). Gene regulatory landscape of cerebral cortex folding. Sci Adv, 10(23), eadn1640. 10.1126/sciadv.adn1640

Sousa, A. M. M., Meyer, K. A., Santpere, G., Gulden, F. O., & Sestan, N. (2017). Evolution of the Human Nervous System Function, Structure, and Development. Cell, 170(2), 226–247. 10.1016/j.cell.2017.06.036

Stahl, R., Walcher, T., De Juan Romero, C., Pilz, G. A., Cappello, S., Irmler, M., Sanz-Aquela, J. M., Beckers, J., Blum, R., Borrell, V., & Götz, M. (2013). Trnp1 regulates expansion and folding of the mammalian cerebral cortex by control of radial glial fate. Cell, 153(3), 535–549. 10.1016/j.cell.2013.03.027

Storbeck, M., Hupperich, K., Gaspar, J. A., Meganathan, K., Martínez Carrera, L., Wirth, R., Sachinidis, A., & Wirth, B. (2014). Neuronal-specific deficiency of the splicing factor Tra2b causes apoptosis in neurogenic areas of the developing mouse brain. PLoS One, 9(2), e89020. 10.1371/journal.pone.0089020

Tirosh, I., Izar, B., Prakadan, S. M., Wadsworth, M. H., 2nd, Treacy, D., Trombetta, J. J., Rotem, A., Rodman, C., Lian, C., Murphy, G., Fallahi-Sichani, M., Dutton-Regester, K., Lin, J. R., Cohen, O., Shah, P., Lu, D., Genshaft, A. S., Hughes, T. K., Ziegler, C. G., … Garraway, L. A. (2016). Dissecting the multicellular ecosystem of metastatic melanoma by single-cell RNA-seq. Science, 352(6282), 189–196. 10.1126/science.aad0501

Tyagi, G., Carnes, K., Morrow, C., Kostereva, N. V., Ekman, G. C., Meling, D. D., Hostetler, C., Griswold, M., Murphy, K. M., Hess, R. A., Hofmann, M. C., & Cooke, P. S. (2009). Loss of Etv5 decreases proliferation and RET levels in neonatal mouse testicular germ cells and causes an abnormal first wave of spermatogenesis. Biol Reprod, 81(2), 258–266. 10.1095/biolreprod.108.075200

Uebbing, S., Gockley, J., Reilly, S. K., Kocher, A. A., Geller, E., Gandotra, N., Scharfe, C., Cotney, J., & Noonan, J. P. (2021). Massively parallel discovery of human-specific substitutions that alter enhancer activity. Proc Natl Acad Sci U S A, 118(2). 10.1073/pnas.2007049118

Uren, P. J., Bahrami-Samani, E., Burns, S. C., Qiao, M., Karginov, F. V., Hodges, E., Hannon, G. J., Sanford, J. R., Penalva, L. O., & Smith, A. D. (2012). Site identification in high-throughput RNA-protein interaction data. Bioinformatics, 28(23), 3013–3020. 10.1093/bioinformatics/bts569

Uzquiano, A., Kedaigle, A. J., Pigoni, M., Paulsen, B., Adiconis, X., Kim, K., Faits, T., Nagaraja, S., Antón-Bolaños, N., Gerhardinger, C., Tucewicz, A., Murray, E., Jin, X., Buenrostro, J., Chen, F., Velasco, S., Regev, A., Levin, J. Z., & Arlotta, P. (2022). Proper acquisition of cell class identity in organoids allows definition of fate specification programs of the human cerebral cortex. Cell, 185(20), 3770–3788.e3727. 10.1016/j.cell.2022.09.010

Velasco, S., Kedaigle, A. J., Simmons, S. K., Nash, A., Rocha, M., Quadrato, G., Paulsen, B., Nguyen, L., Adiconis, X., Regev, A., Levin, J. Z., & Arlotta, P. (2019). Individual brain organoids reproducibly form cell diversity of the human cerebral cortex. Nature, 570(7762), 523–527. 10.1038/s41586-019-1289-x

Walter, C., Balouchzadeh, R., Garcia, K. E., Kroenke, C. D., Pathak, A., & Bayly, P. V. (2023). Multi-scale measurement of stiffness in the developing ferret brain. Sci Rep, 13(1), 20583. 10.1038/s41598-023-47900-4

Wang, L., Hou, S., & Han, Y. G. (2016). Hedgehog signaling promotes basal progenitor expansion and the growth and folding of the neocortex. Nat Neurosci, 19(7), 888–896. 10.1038/nn.4307

Wei, Y., Han, S., Wen, J., Liao, J., Liang, J., Yu, J., Chen, X., Xiang, S., Huang, Z., & Zhang, B. (2023). E26 transformation-specific transcription variant 5 in development and cancer: modification, regulation and function. J Biomed Sci, 30(1), 17. 10.1186/s12929-023-00909-3

Whalen, S., Inoue, F., Ryu, H., Fair, T., Markenscoff-Papadimitriou, E., Keough, K., Kircher, M., Martin, B., Alvarado, B., Elor, O., Laboy Cintron, D., Williams, A., Hassan Samee, M. A., Thomas, S., Krencik, R., Ullian, E. M., Kriegstein, A., Rubenstein, J. L., Shendure, J., … Pollard, K. S. (2023). Machine learning dissection of human accelerated regions in primate neurodevelopment. Neuron, 111(6), 857–873.e858. 10.1016/j.neuron.2022.12.026

Won, H., de la Torre-Ubieta, L., Stein, J. L., Parikshak, N. N., Huang, J., Opland, C. K., Gandal, M. J., Sutton, G. J., Hormozdiari, F., Lu, D., Lee, C., Eskin, E., Voineagu, I., Ernst, J., & Geschwind, D. H. (2016). Chromosome conformation elucidates regulatory relationships in developing human brain. Nature, 538(7626), 523–527. 10.1038/nature19847

Won, H., Huang, J., Opland, C. K., Hartl, C. L., & Geschwind, D. H. (2019). Human evolved regulatory elements modulate genes involved in cortical expansion and neurodevelopmental disease susceptibility. Nat Commun, 10(1), 2396. 10.1038/s41467-019-10248-3

Xu, K., Schadt, E. E., Pollard, K. S., Roussos, P., & Dudley, J. T. (2015). Genomic and network patterns of schizophrenia genetic variation in human evolutionary accelerated regions. Mol Biol Evol, 32(5), 1148–1160. 10.1093/molbev/msv031

Yabut, O. R., Fernandez, G., Huynh, T., Yoon, K., & Pleasure, S. J. (2015). Suppressor of Fused Is Critical for Maintenance of Neuronal Progenitor Identity during Corticogenesis. Cell Rep, 12(12), 2021–2034. 10.1016/j.celrep.2015.08.031

Zhang, X., Zhao, X., Li, G., Zhang, M., Xing, P., Li, Z., Chen, B., Yang, H., & Wu, Z. (2021). Establishment of Etv5 gene knockout mice as a recipient model for spermatogonial stem cell transplantation. Biol Open, 10(1). 10.1242/bio.056804

Zhao, J., Li, D., Seo, J., Allen, A. S., & Gordân, R. (2017). Quantifying the Impact of Non-coding Variants on Transcription Factor-DNA Binding. Res Comput Mol Biol, 10229, 336–352. 10.1007/978-3-319-56970-3_21

Zhu, Y., Sousa, A. M. M., Gao, T., Skarica, M., Li, M., Santpere, G., Esteller-Cucala, P., Juan, D., Ferrández-Peral, L., Gulden, F. O., Yang, M., Miller, D. J., Marques-Bonet, T., Imamura Kawasawa, Y., Zhao, H., & Sestan, N. (2018). Spatiotemporal transcriptomic divergence across human and macaque brain development. Science, 362(6420). 10.1126/science.aat8077

